# A corset of adhesions during development establishes individual neural stem cell niches and controls adult behaviour

**DOI:** 10.1101/2022.10.04.510893

**Authors:** Agata Banach-Latapy, Vincent Rincheval, David Briand, Isabelle Guénal, Pauline Spéder

## Abstract

Neural stem cells (NSCs) reside in a defined cellular microenvironment, the niche, which supports the generation and integration of neuronal lineages. The mechanisms building a sophisticated niche structure around NSCs, and their functional relevance for neurogenesis are yet to be understood. In the *Drosophila* larval brain, the cortex glia (CG) encase individual NSC lineages, organizing the stem cell population and newborn neurons into a stereotypic structure. We first found that lineage information is dominant over stem cell fate. We then discovered that, in addition to timing, the balance between multiple adhesion complexes supports the individual encasing of NSC lineages. An intra-lineage adhesion through homophilic Neuroglian interactions provides strong binding between cells of a same lineage, while a weaker interaction through Neurexin-IV exists between CG to NSC lineages. Their loss leads to random, aberrant grouping of several NSC lineages together, and to altered axonal projection of newborn neurons. Further, we link the loss of these two adhesion complexes during development to locomotor hyperactivity in the resulting adults. Altogether, our findings identify a corset of adhesions building a neurogenic niche at the scale of individual stem cell and provide the proof-of-principle that mechanisms supporting niche formation during development define adult behaviour.

## INTRODUCTION

Stem cells are multipotent progenitors driving the growth and regeneration of the tissue they reside in through the generation of differentiated cells. Their localisation within the tissue is restricted to carefully arranged cellular microenvironments, or niches, which control their maintenance and activity in response to local and systemic cues [1–3]. The niches comprise the stem cell themselves, their newborn progeny and a number of cells of various origins and roles that support stem cell decisions. The diversity of cellular shapes and roles requires a precise spatial organisation to enable proper niche function towards all and every stem cells. Within the central nervous system (CNS) in particular, a highly structured organ dependent on the tight arrangement of cellular connections, the neural stem cell (NSC) niches are anatomically complex microenvironments that must form within such constraint. They comprise multiple cell types such as neurons, various glial cells, vasculature and immune cells [4,5] which are precisely organised with respect to NSCs. While studies have focused on the identification of signaling pathways operating in an established niche and controlling neurogenesis [5–7], how the niche is first spatially built around NSCs, and the importance of its architecture on neurogenesis, from stem cell division to the integration of the newborn neurons, are poorly understood.

The *Drosophila* larval CNS offers a genetically powerful model to study interactions within the niche *in vivo*. Similar to mammals, *Drosophila* NSCs, historically called neuroblasts, self-renew to produce neuronal and glial progeny, and their behaviour is controlled by their niche, an exquisitely organised yet less complex structure than its mammalian counterpart.

Fly NSCs are born during embryogenesis, during which they cycle to generate primary neurons in a first wave of neurogenesis. They then enter quiescence, a mitotically dormant phase from which they exit to proliferate through the autonomous activation of PI3K/Akt in response to nutrition [8,9]. This post-embryonic, second wave of neurogenesis generates secondary neurons that will make up 90 % of the adult CNS and lasts until the end of the larval stage. NSCs finally differentiate or die by apoptosis after pupariation. Larval NSCs populate the different regions of the CNS, namely the ventral nerve cord (VNC), the central brain (CB) and the optic lobe (OL) (Fig. 1A). They nevertheless display distinct properties, mainly through different modes of division and expression of specific transcription factors (Fig. 1B) [10]. Type I NSCs, the most represented subtype, reside in the CB and VNC, and divide asymmetrically to generate a smaller ganglion mother cell (GMC). GMCs further terminally divide to produce two neurons. Type II NSCs, found exclusively in the CB, represent a smaller population with 8 cells per brain hemisphere [11–13]. Type II NSC self-renewal produces an Intermediate Neural Progenitor (iINP), which undergoes a limited number of asymmetric divisions to produce GMCs that will subsequently divide to give neurons.

**Figure 1.**
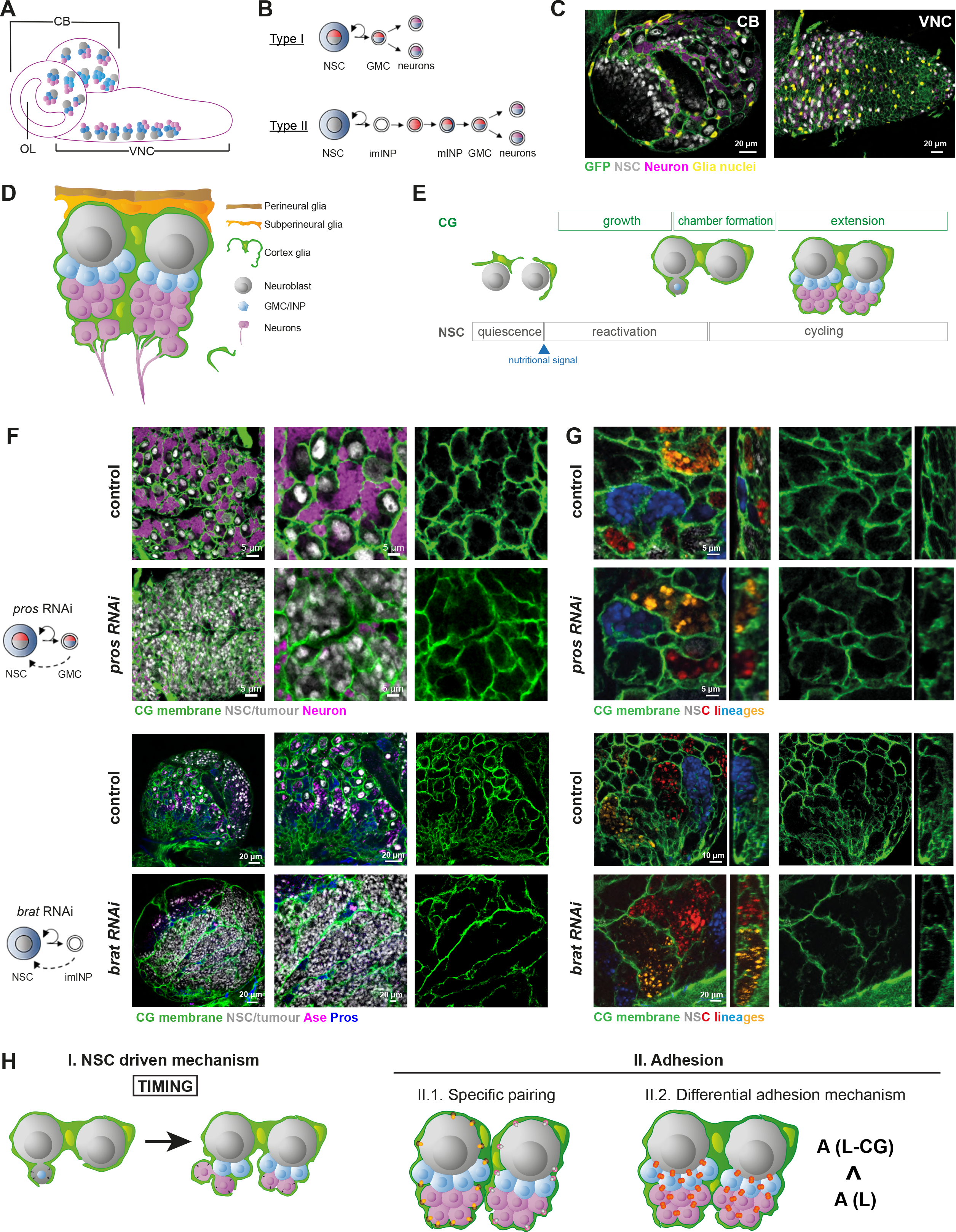
Cortex glia recognize lineage over identity to generate stereotypic encasing of NSC lineages. A) Schematic of the Drosophila larval CNS depicting the localisation of the NSC lineages. Two main neurogenic regions are the central brain (CB), comprising two hemispheres, and the ventral nerve cord (VNC). B) Schematic of the Type I and Type II NSC lineages. C) Confocal pictures representing the dorsal region of the larval CB and ventral region of the VNC at ALH72 (at 25°C) labelled with markers for the CG membranes (*Nrv2∷GFP*, green), glia nuclei (anti-Repo, yellow), NSC (anti-Dpn, grey) and neurons (anti-ElaV, magenta). D) Schematic of the NSC niche, made by the perineurial glia (PG, brown), subperineurial glia (SPG, orange), cortex glia (CG, green), neural stem cells (NSC, grey), ganglion mother cells/intermediate progenitors (gmc/inp, blue) and neurons (N, magenta). E) Timeline of the encasing of NSC lineages by CG parallel to NSC behaviour. F) Adaptation of the CG network to NSC tumours. Type I (*wor-GAL4 > pros RNAi*) and Type II (*PntP1-GAL4 > brat RNAi*) tumours were induced and the organisation of CG membrane was monitored by *Nrv2∷GFP* (green). NSCs are labelled with Dpn (grey), and neuron with ElaV (magenta). See Methods for timing, conditions and genetics of larval rearing. G) Relationship between individual encasing and cell identity. Type I (*wor-GAL4 > pros RNAi*) and Type II (*PntP1-GAL4 > brat RNAi*) tumours were induced together with the multicolour lineage tracing Raeppli-NLS. One out of four colours (blue, white, orange and red) is stochastically chosen in the transformed NSC upon induction Heat shock induction for 2 h at 37°C was performed at ALH0 using *hs-Flp*. CG membrane is visualized with *Nrv2∷GFP* (green). See Methods for timing, conditions and genetics of larval rearing. H) Schematic of the different hypotheses explaining individual encasing of NSC lineages by CG. Panel I depicts the NSC-driven timing of NSC encapsulation, prior to lineage generation, as the instructive cue. Panel II describes the use of adhesion mechanisms: 1. Specific CG to NSC lineage and 2. Generic, based on difference in strength between intra-lineage adhesion (A_L_) and adhesion linking NSC lineages and CG (A_L-CG_); in this case, the prediction is that A_L_ > A_L-CG_.

These different NSCs are embedded within a sophisticated, multi-layered niche made of different cell types (Fig. 1C-D). The blood-brain barrier forms the interface with the systemic environment, and controls NSC reactivation [14] and proliferation [15]. A specific glial subtype, the cortex glia (CG), is in close contact with NSCs and their progeny, and is crucial for NSC proliferation [16–18], resistance to stress [19,20] as well as for the survival of newborn neurons [21]. Remarkably, the CG form a continuous glial network which invades the whole CNS (Fig 1C-D) while building bespoke encasing of individual NSC lineages (comprising NSC, GMC and newborn neurons, as well as INP for Type II NSCs), called CG chambers [21–23]. CG also enwrap in individual encasing primary neurons and, later on, older, mature secondary neurons (Fig. 1D). CG network is progressively built around NSC during larval development in a process which parallels NSC behaviour [21,24,25] (Fig.1E). CG cells, born during embryogenesis, do not form a continuous meshwork nor encase quiescent NSCs at larval hatching. Rather, they initiate growth in response to nutrition, via autonomous activation of the PI3K/Akt pathway, leading to an increase in membrane density yet without NSC encasing. Then, at the time NSCs start dividing, CG enwrap individual NSCs, forming a typical chequerboard structure. They further extend their processes to maintain a fitted chamber structure during neuronal production. The cell bodies of newborn neurons from one NSC lineage are thus initially found clustered together in one CG chamber; as they mature, they will be individually encased by the CG. Newborn, still immature neurons from the same lineage start to extend axonal projections, which are fasciculated together as a bundle and are also encased by the CG (Fig. 1D and Supp. Fig. 1A-A’) until they enter the neuropile, a synaptically dense region devoided of cell bodies, where axons connect [26–28]. There, they will be taken care of by other glial cell types [29]. The repeated pattern of individual chamber thus translates both in term of cell bodies and axonal tracts.

The reliable formation of such precise chequerboard structure implies that CG integrate proper cellular cues to decide whether to encase specific cells, while navigating between a density of diverse cell types. However, the nature of these cues, and the importance of such stereotypicencasing of NSC lineages on NSC activity and generation of functional neuronal progeny remained to be identified. We first found that CG are able to distinguish between lineages and that lineage information prevails over cell identity. Further, we discovered that lineage information and individual encasing are mediated by the existence of multiple adhesion complexes within the niche. First, the cell adhesion protein Neurexin-IV is expressed and crucial in NSC lineages to maintain their individual encasing, through its interaction with Wrapper, a protein with immunoglobulin domain present in the CG. In parallel, Neuroglian appears to form strong homophilic interactions between cells of the same lineages, keeping them together by providing a stronger adhesion compared to the weaker interaction between CG and NSC lineages. In absence of either Neurexin-IV and Neuroglian, NSC lineages are grouped in a random fashion. The loss of these adhesions is further associated with misprojected axonal bundles. Adherens junctions are also present in NSC lineages, however they appear mostly dispensible for indvidual encasing. Further, we demonstrated that the loss of Neurexin-IV and Neurglian adhesions during development is linked to an altered, hyperactive locomotor behaviour in the adult. Our findings unravel a principle of NSC niche organization based on a differential in adhesion and link the adhesive property of the niche during development to adult neurological behaviour.

## RESULTS

### Cortex glia distinguish between NSC lineages

Before lineage generation, CG encase only one, and not several NSCs within a membranous chamber, suggesting that CG can sense cell identity to decide which cell to enwrap. To assess the importance of NSC fate in chamber formation, we took advantage of genetic alterations known to dysregulate NSC division and differentiation and lead to the formation of tumour-like, NSC-only, lineages [30]. In particular, *pros* knockdown in Type I lineages converts GMC into NSC-like, Dpn^+^ cells at the expense of neurons [31]. Surprisingly, in these conditions, we found that CG chambers contained not one, but several NSC-like, Dpn^+^ cells (Fig. 1F). Similar results were obtained for other conditions that lead to Type I NSC-only lineages, including GMC dedifferentiation via Dpn overexpression or loss of asymmetric division via aPKC overexpression (Supp. Fig. 1B). We then asked how CG would adapt to the dysregulation of another type of NSCs, the Type II NSC. Since CG chamber formation was precisely described only for type I NSCs [21], we first checked the dynamics of CG morphogenesis around type II and found that they followed similar steps, albeit in a slower fashion (Supp. Fig. 1C). We then knocked down the cell fate determinant *brat* [32,33], which is necessary for preventing iINP dedifferentiation into NSC-like cells. This led to the formation of large tumours of Type II NSCs (Fig 1F). CG were able to adapt to the outgrowth of cells and enwrapped many Type II NSCs within one chamber. These data show that both for Type I and Type II NSCs, stem cell identity is not sufficient to ensure their sorting into individual CG chambers.

We then wondered whether tumour NSCs grouped within one chamber originated from the same NSC mother cell or had been encased randomly independently of their lineage of origin. To do so, we used multi-colour clonal analysis to label individual NSC lineages. The Raeppli system [34] allows the stochastic and irreversible labelling of a cell at the time of induction, allowing to mark and track a mother cell and its colour-sharing progeny. Induction before NSC reactivation of a nuclear tagged version of Raeppli (Raeppli-NLS) ensured all lineages could be fully tracked. We first confirmed that cells found within each CG chamber belonged to the same lineage for wild-type Type I and Type II NSC lineages (Fig 1G). We then found that clonal tumour-like growth coming from single dysregulated NSCs were contained within one CG chamber, both for Type I and Type II NSCs (Fig. 1G). Altogether, these results demonstrate that CG are able to distinguish between different NSC lineages, and that keeping a lineage together prevails over encasing individual NSC, thus showing that the lineage information prevails over cell identity.

### Individual encasing relies on intrinsic lineage cues

We then assessed the molecular basis of NSC lineage recognition and individual encasing by CG. Chamber completion around NSCs occurs around the time of first division and is driven by NSC reactivation [21,24]. A simple explanation for keeping a lineage together and separated from others would thus be the sequential addition of newborn cells within a compartment already defined from the start, by NSC-derived signals (Fig.1H, Panel I). Timing would thus be the instructive cue. Indeed, previous studies have shown that instructions from reactivated NSCs are paramount to form CG niches [21], although the cues integrated by the glia have not been identified. Within this hypothesis, blocking CG morphogenesis until well after the first NSC division, followed by subsequent release of their growth, would result in aberrant chamber formation and random encasing of neurons from different lineages.

To do so, we prevented chamber formation by blocking the activation of the PI3K/Akt pathway (PTEN inhibitor) specifically in the CG [21], using the QF system to control the timing of induction (Fig. 2A). At the same time, NSC lineages were stochastically marked using Raeppli-NLS, driven this time by the GAL4/UAS system. CG growth was prevented until NSCs cycled actively and neuronal progeny had been already produced, resulting in CG failing to form correct chambers to separate individual lineages from each other (Supp. Fig. 2A, time T1). We then allowed CG to resume their growth and observed the establishment of a stereotyped chequerboard pattern, with most of the chambers containing single NSC and neurons (Fig. 2B, time T2). This was confirmed by quantifying the chambers containing more than on NSC lineage (Fig. 2C). These results thus show that the information allowing CG to distinguish between lineages is not an exclusive property of the NSC, but is likely inherited by their progeny, and can be sensed by CG later during larval development.

**Figure 2.**
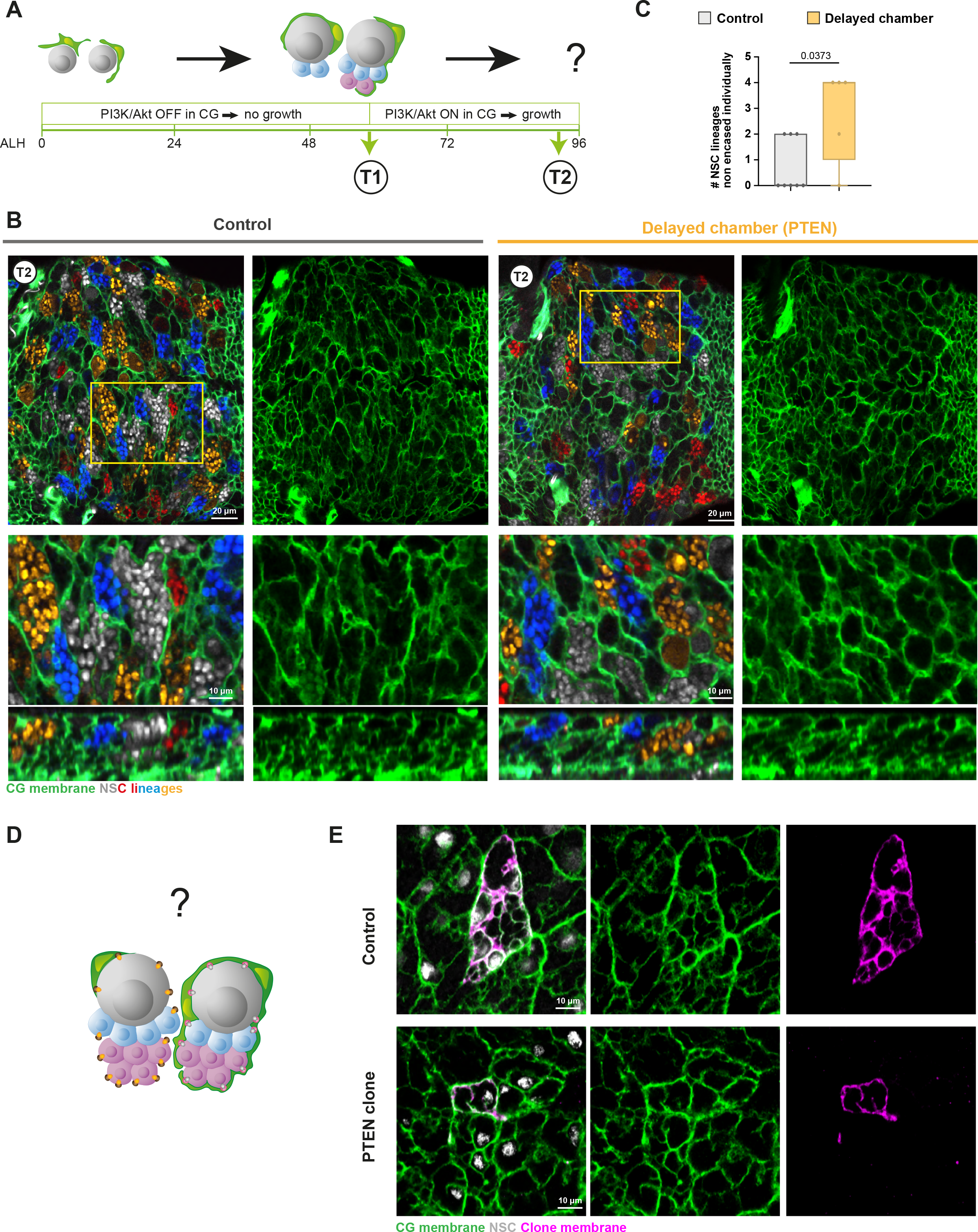
Cortex glia use intrinsic and generic NSC lineage cues for individual encasing. A) Schematic of the timing and genetic conditions used to probe the importance of NSC-driven timing of NSC encapsulation in keeping NSC lineages together. CG growth is initially blocked using PTEN expression in the CG. At T1, the block is removed, and CG structure is assessed at T2. B) Representative confocal picture of the extent of individual encasing of NSC lineages by CG after the regimen described in A. Top panel shows the whole thoracic VNC, and bottom panel a close-up of the yellow box. NSC lineages were marked with the multicolour lineage tracing Raeppli-NLS (blue, white, orange and red), induced at ALH0 using *hs-Flp*. CG membrane is visualized with *Nrv2∷GFP* (green). See Methods for timing, conditions and genetics of larval rearing. C) Quantification of the number of NSC lineages non-individually encased from B). Control (n = 8 VNCs) and PTEN conditional block (n = 5 VNCs). Data statistics: Mann-Whitney test. Results are presented as box and whisker plots. D) Schematic of the experiment designed to probe whether specific, non-interchangeable adhesions exist between individual CG cells and individual NSC lineages. E) Representative confocal picture of the CG network in control CG clone and in clone in which CG growth was blocked (PTEN). The CoinFLP system was used to generate wild-type and PTEN clones in the CG. See Methods for timing, conditions and genetics of larval rearing. The membrane of the clone is marked by mCD8∷RFP (magenta) CG membrane is visualized with *Nrv2∷GFP* (green).

One elegant way cells could be recognised by CG as belonging to the same lineage would be to keep them physically together through adhesion mechanisms. We hypothesized two possibilities (Fig. 1H, Panel II).

The most complex mechanism would rely on the existence of lineage-specific adhesions, with a code of unique molecular interactions between specific CG cells and all cells of specific lineages (Fig. 1H, Panel II.1). In this context, CG cells would not be interchangeable.

The simplest solution would see all NSC lineages using the same adhesion mechanisms to link their cells (Fig. 1H, Panel II.2). The differential adhesion hypothesis proposes that cells with similar adhesive strength cluster together, ultimately sorting cell populations with different adhesions and creating cellular compartments [35]. While NSC lineages and CG are not fully sorted in only two compartments, NSC encapsulation could be seen as a local segregation event between whole NSC lineages and the CG. NSC lineages with intra-lineage adhesions (A_L_) stronger than their adhesion to the CG (A_L-CG_) would form a physical barrier for the CG, preventing their intercalation in between cells from the same lineage and thus leading to encapsulation of the whole lineage. In this case, a difference in adhesive properties would be sufficient to segregate CG from NSC lineages, and CG cells would be interchangeable.

To discriminate between these two hypotheses, we first assessed the result of preventing some CG cells to cover NSC lineages to a normal extent (Fig. 2D). We used clonal analysis to randomly impair the growth of a few CG cells within the entire population, through a Coin-FLP approach [36]. Blocking PI3K/Akt signaling in a few cells (marked with RFP) through the expression of the inhibitor PTEN resulted in much smaller clones compared to control wild-type clones (Fig. 2E). However, the CG network itself appeared gapless around PTEN clones, revealing that CG were able to compensate for the loss of their neighbours’ membrane to restore NSC chambers (Fig. 2E). To confirm that CG cells are completely interchangeable we used the same approach to kill a few CG cells, expressing the pro-apoptotic gene *reaper* [37] to induce apoptosis only once the NSC chambers were already formed (Supp. Fig 2B). Induction of apoptosis led to a near complete loss of RFP-marked clones, only visible through cell remnants. Nevertheless, the whole CG network appeared intact. This shows that any given NSC lineage can recruit other CG cells, and that, in accordance with our previous findings (Fig 2A-C), it is able to do so after NSC reactivation. Similar results demonstrating the ability of CG to replace each other has been previously obtained around neurons [38]. Altogether, our results indicate that NSC lineages can be enwrapped by different CG already taking care of other lineages, and that specific CG-NSC lineage pairings do not occur. This suggests that the same adhesion mechanism might be repeated for each NSC lineage, providing stronger cohesion between cells of the same NSC lineage than between CG and NSC lineages.

### Intra-lineage adherens junctions are present but not absolutely required for encasing of individual NSC lineages

Previous studies had reported the localization within larval NSC lineages of the Drosophila E-cadherin Shotgun (Shg), a component of adherens junctions usually present in epithelia [22,39]. We analysed the expression pattern of Shg during larval development, using a protein trap fusion (*Shg∷GFP*, Fig. 3A). Shg∷GFP was detected from larval hatching, initially present around and between NSCs (ALH24, dashed yellow circle), and also along CG membranes (white arrowheads). Shg∷GFP was then no longer detected between NSCs having proceeded through reactivation, yet not individually encased (ALH48, yellow stars). At later larval stages, following neuronal production, Shg∷GFP showed a remarkable pattern of expression, with a strong enrichment between cells from the same lineage (ALH72). A similar pattern was found for its ß-catenin partner Armadillo (Fig. 3B) through antibody staining. As Shg form homophilic bonds, this suggests that adherens junctions exist between cells of the same NSC lineage.

**Figure 3.**
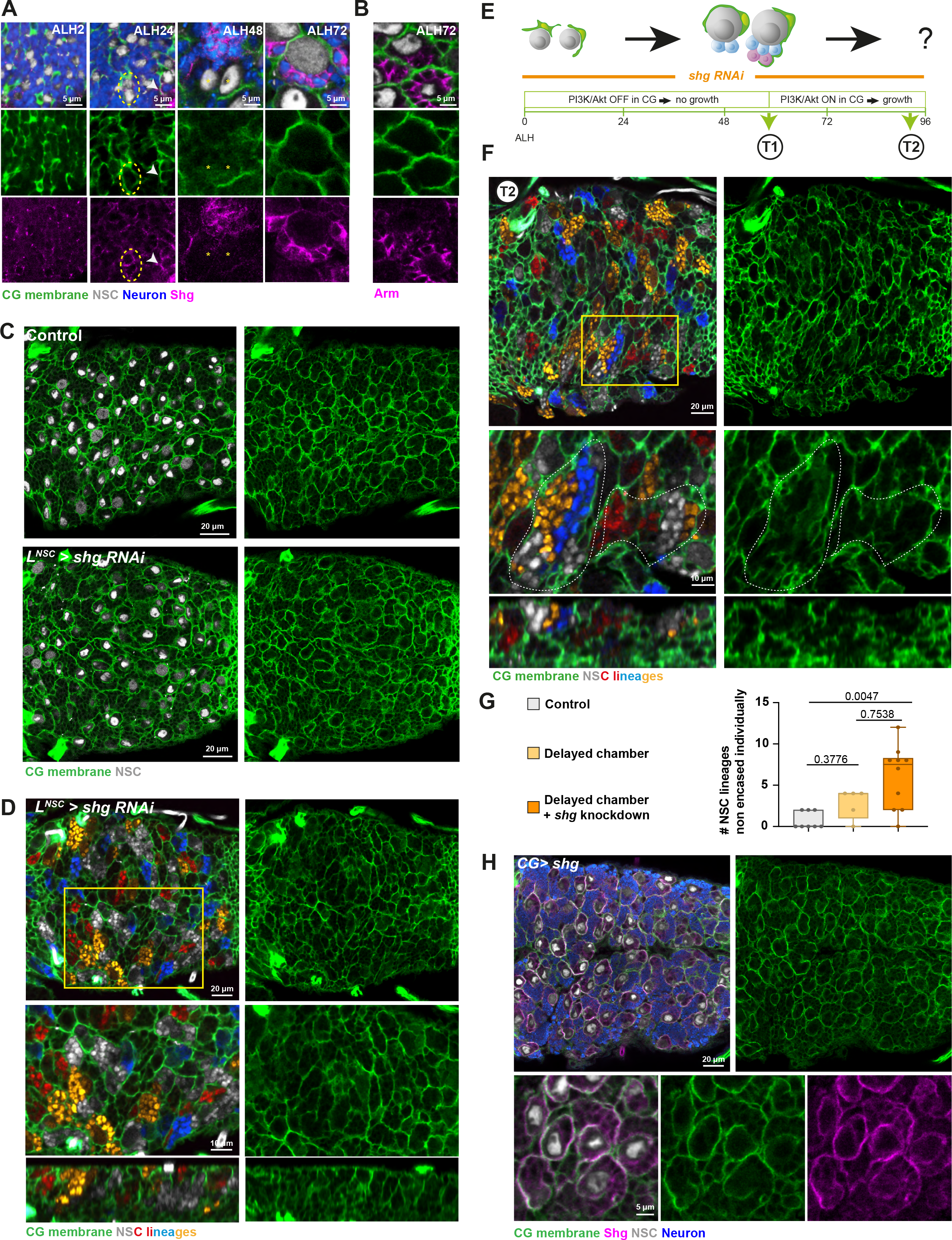
Intra-lineage adherens junctions are not absolutely required for individual encasing while providing robustness. A) Representative confocal images of the expression of the *Drosophila* E-cadherin, Shg, at ALH0, ALH24, ALH48 and ALH72 at 25°C. Shg is monitored through a *Shg∷GFP* fusion (magenta), CG membrane is visualized by *Nrv2∷GFP* (green), NSCs are labelled with Dpn (grey), and neurons are labelled with ElaV (blue). B) Representative confocal images of the expression of the *Drosophila* β-catenin, Arm, at ALH72 at 25°C. Arm is detected with a specific antibody (magenta), CG membrane is visualized by *Nrv2∷GFP* (green), NSCs are labelled with Dpn (grey), and neurons are labelled with ElaV (blue). C) Representative confocal pictures of the thoracic VNC for control and *shg* knockdown by RNAi in NSC lineages (*L^NSC^ > shg RNAi*, driver line *Nrv2∷GFP, wor-GAL4; tub-GAL80^ts^*). Larvae are dissected after 68 h at 29°C from ALH0. CG membrane is visualized by *Nrv2∷GFP* (green) and NSCs are labelled with Dpn (grey). D) Representative confocal picture of the thoracic VNC for *shg* knockdown by RNAi in NSCs marked with the multicolour lineage tracing Raeppli-NLS (blue, white, orange and red-. Raeppli-NLS is induced at ALH0 using *hs-Flp*. Larvae are dissected after 68 h at 29°C from ALH0. CG membrane is visualized with *Nrv2∷GFP* (green). See Methods for timing, conditions and genetics of larval rearing. E) Schematic of the timing and genetic conditions used to probe the importance of Shg adhesion (*shg* RNAi) on individual encasing of NSC lineages when CG growth is initially blocked (PTEN). At T1, the PTEN block is removed, and CG structure is assessed at T2. F) Representative confocal picture of the extent of individual encasing of NSC lineages by CG after the regimen described in E). Top panel shows the whole thoracic VNC, and bottom panel a close-up of the yellow box. Dashed white lines highlight NSC clones encased together. NSC lineages were marked with the multicolour lineage tracing Raeppli-NLS (blue, white, orange and red), induced at ALH0 using *hs-Flp*. CG membrane was visualized with *Nrv2∷GFP* (green). See Methods for timing, conditions and genetics of larval rearing. G) Quantification of the number of NSC lineages non-individually encased from F). Control (n = 8 VNCs), PTEN conditional block (n = 5 VNCs) and PTEN conditional block + *shg* RNAi (n = 10 VNCs). Data statistics: one-way ANOVA with a Kruskal– Wallis multiple comparison test. Results are presented as box and whisker plots. H) Representative confocal picture of the thoracic VNC and close-up for *shg* overexpression in the CG (driver *Nrv2∷GFP, tub-GAL80^ts^; cyp4g15-GAL4*). Larvae are dissected after 68 h at 29°C from ALH0. CG membrane is visualized by *Nrv2∷GFP* (green), NSCs are labelled with Dpn (grey), neurons are labelled with ElaV (blue) and Shg is detected with a specific antibody (magenta).

We then wondered whether such adhesion was a property of differentiating lineages within the CG chamber, or whether it could take place between cells of similar fate and identity. We first looked at NSC-like cells from *pros* and *brat* tumours, which we showed are contained clonally within one CG chamber (Fig. 1G). A strong Shg staining was detected between NSC-like cells from the same lineage (Supp. Fig. 3A). It was however not the case when they originated from different lineages, which were separated by CG membrane. This suggests that adherens junctions are not only a property of differentiating lineage, but, importantly, is found between cells in an inverse correlation with the presence of CG membranes. In line with this finding, we observed a strong Shg staining between NSCs of the optic lobe, another type of neural progenitors [40] which are contained within one CG chamber (Supp. Fig. 3B). Altogether these results suggest that AJ could be a mean of keeping lineages together, providing differential adhesion that would ensure stronger bond between cells of the same lineage than between lineages and CG, and functioning as a physical barrier to further enwrapping of individual cells by CG membranes.

We first asked whether adherens junctions were necessary for keeping cells together within the CG chamber. Previous studies indeed suggested that E-cadherin expression was required in NSC lineages for proper CG structure [22,41]. We first generated NSC lineages mutant for *shg*, by inducing MARCM clones during late embryogenesis (see Methods). To our surprise, Shg-depleted NSC lineages still stayed individually encased within one CG chamber (Supp. Fig. 3C, upper panel). The same result was obtained when clones were induced at a later timepoint, to prevent potential compensation through the upregulation of other adhesion molecules (Supp. Fig. 3C, lower panel). In accordance with these results, we found that RNAi knockdown of *shg* in NSC or neurons did not disrupt overall lineage organization within CG chambers (Fig. 3C), despite successfully removing Shg∷GFP signal (Supp. Fig. 3D). To note, we also did not record disruption of CG network when *shg* was knocked down in CG themselves (Supp. Fig 3E), contrary to previous findings [22]. Driving the multicolour clonal marker Raeppli-NLS (induced at ALH0) along with *shg* RNAi driven from embryogenesis in the NSC lineage demonstrated the conservation of individual encasing despite efficient E-cadherin loss (Fig. 3D and Supp. Fig. 3F-G). These results argue against the strict requirement of Shg-mediated adhesion for NSC lineage maintenance within one CG chamber [22,41]. All together, these data suggest that adherens junctions, while expressed within the NSC lineage, are not absolutely required for their individual encasing.

We then wondered whether intra-lineage AJ could rather be used as a safety mechanism, ensuring robustness in a system in which other strategies would primarily provide intra-lineage cohesion. As the CG chamber encases NSC at the time they produce their first progeny, timing would bring a first level of clustering (Fig. 1H, case I), while AJ would ensure its maintenance in the case the chamber is affected (such as in Fig. 2A-C). To test this hypothesis, we performed a conditional block of CG growth while at the same time constantly driving *shg* RNAi and the multicolour clonal tool Raeppli-NLS in NSC lineages (Fig. 3E and Supp. Fig. 3H). The efficiency of *shg* knockdown was assessed through E-cadherin staining (Supp. Fig. 3F). In this case, the progeny born after the re-establishment of CG growth will naturally be encased, while before will depend on the requirement of adherens junctions in absence of proper encasing. Looking at the slightly deeper level in which differentiating progeny reside, we uncovered few restricted, localised defects in the individual encasing of NSC lineages, with several colours detected within the boundaries of one continuous CG membrane (Fig. 3F-G). However, most lineages still appear encased correctly These results indicate that adherens junctions might participate in the robustness of individual NSC lineage encasing when CG are altered, however in a limited fashion.

Finally, we wondered whether altering the respective adhesion balance between NSC lineages and CG would shift the site of preferential adhesion, and thus alter the sorting of the different cell types. Ultimately, this would result in randomizing the number of NSC lineage encased within one CG chamber. To do so, we overexpressed *shg* in the CG, with the aim to force the recruitment of lineage-expressed endogenous Shg to adherens junctions artificially set up between CG and NSC lineage cells (Fig. 3H), and as such to flatten the E-cadherin based adhesion difference (with now A_L_ = A_L-CG_ for Shg). Lineage-expressed endogenous Shg would thus have the choice to generate adhesion of similar strength either with itself, or with CG-provided Shg. Despite the successful expression of *shg* in the CG, the usual pattern of CG chambers was nevertheless maintained in this condition (Fig. 3H), suggesting that in this case A_L_ still stays superior to A_L-CG_, and that other adhesions exist to fulfill this role. These data imply that a difference in adhesion using adherens junction is not the main driver for ensuring the individual encasing of NSC lineages.

### Occluding junction components are expressed in NSC lineages

These findings prompted us to investigate the potential presence and role of other adhesion complexes which could provide intra-lineage cohesion and differential adhesion to sort NSC lineages from CG.

Occluding junctions (tight junctions in vertebrate and septate junctions in *Drosophila*) [42,43] primarily perform a permeability barrier function to paracellular diffusion, mostly described and understood in epithelia or epithelial-like cells. However, they can also provide some adhesion between the cells they link, albeit possibly in a weaker fashion than adherens junctions. *Drosophila* septate junctions are formed by the assembly of cell surface adhesion molecules that can interact in *cis* or *trans*, in an homologous or heterologous fashion, and which are linked to the intracellular milieu by supporting membrane or cytoplasmic molecules [42]. A core, highly conserved tripartite complex of adhesion molecules comprises Neuroglian (Nrg), Contactin (Cont) and Neurexin-IV (Nrx-IV). Nrg, the *Drosophila* homolog of Neurofascin-155, is mostly a homophilic transmembrane protein belonging to the L1-type family and containing several immunoglobulin domains. Cont, homologous to the human Contactin, also contains immunoglobulin domains, is GPI-anchored and only performs heterophilic interactions. Nrx-IV, homologous to the human Caspr/Paranodin, is transmembrane protein with a large extracellular domain including multiple laminin-G domains and EGF repeats [44] and is able to set up heterophilic interactions. In vertebrates, Caspr/Paranodin and Neurofascin-155 are also partners at the paranodal junction between glia and neuron [42]. Besides adhesion molecules, several cytoplasmic or membrane-associated proteins participate in septate junction formation, such as the FERM-family Coracle (Cora), the MAGUK protein Discs large (Dlg1), and the integral membrane Na K-ATPase pump (ATPα).

Interestingly, Nrx-IV and Nrg also perform nervous system-specific roles outside of the septate junction. Nrx-IV is required in the embryonic CNS for the wrapping of individual axon fascicles by midline glia [45–47]. Similarly, neuronal expression of Nrg is important for axonal guidance and regulation of dendritic arborization of peripheral neurons by glial [48,49], and epidermal cells [50], as well as for the function and axon branching of specific larval CB neurons [51,52]. For both proteins, their role in NSC lineages during larval neurogenesis is however poorly known.

We started by assessing the expression and function of septate junction (SJ) components in the larval CNS, and found that Nrx-IV, Nrg, Cora, Dlg1 and ATPα are enriched in NSC lineages.

First, using a protein trap (*Nrx-IV∷GFP*, Fig. 4A), we found that Nrx-IV expression was detected from early larval stage in embryonic (primary) neurons, and around NSCs (ALH0), an expression maintained while NSCs proceed through reactivation (ALH24, dashed yellow circle). As NSCs have reactivated and CG grown (ALH48), Nrx-IV∷GFP appears expressed at the interface between NSC and CG (yellow arrow), but not anymore between NSCs (yellow stars). Further, accompanying the production of newborn, secondary neurons (ALH72), Nrx-IV∷GFP is found enriched at the interface of the cells from the same lineage (NSC, GMC and neurons), while maintaining a strong expression at the interface with CG (yellow arrowhead). A protein trap for Nrg (*Nrg∷GFP*, Fig. 4B) also revealed a strong enrichment between cells of the same NSC lineages following progeny production (ALH72). At this time, Nrg appears enriched in the axonal bundles leaving from secondary neurons, in accordance with previous findings [53]. Moreover, Nrg∷GFP is detected at the interface between lineages and CG (yellow arrow). However, contrary to what we observed with Nrx-IV∷GFP, Nrg∷GFP did not appear enriched between NSCs at any time point before their encasing by CG (ALH0-48, yellow stars and dashed yellow circle).

**Figure 4.**
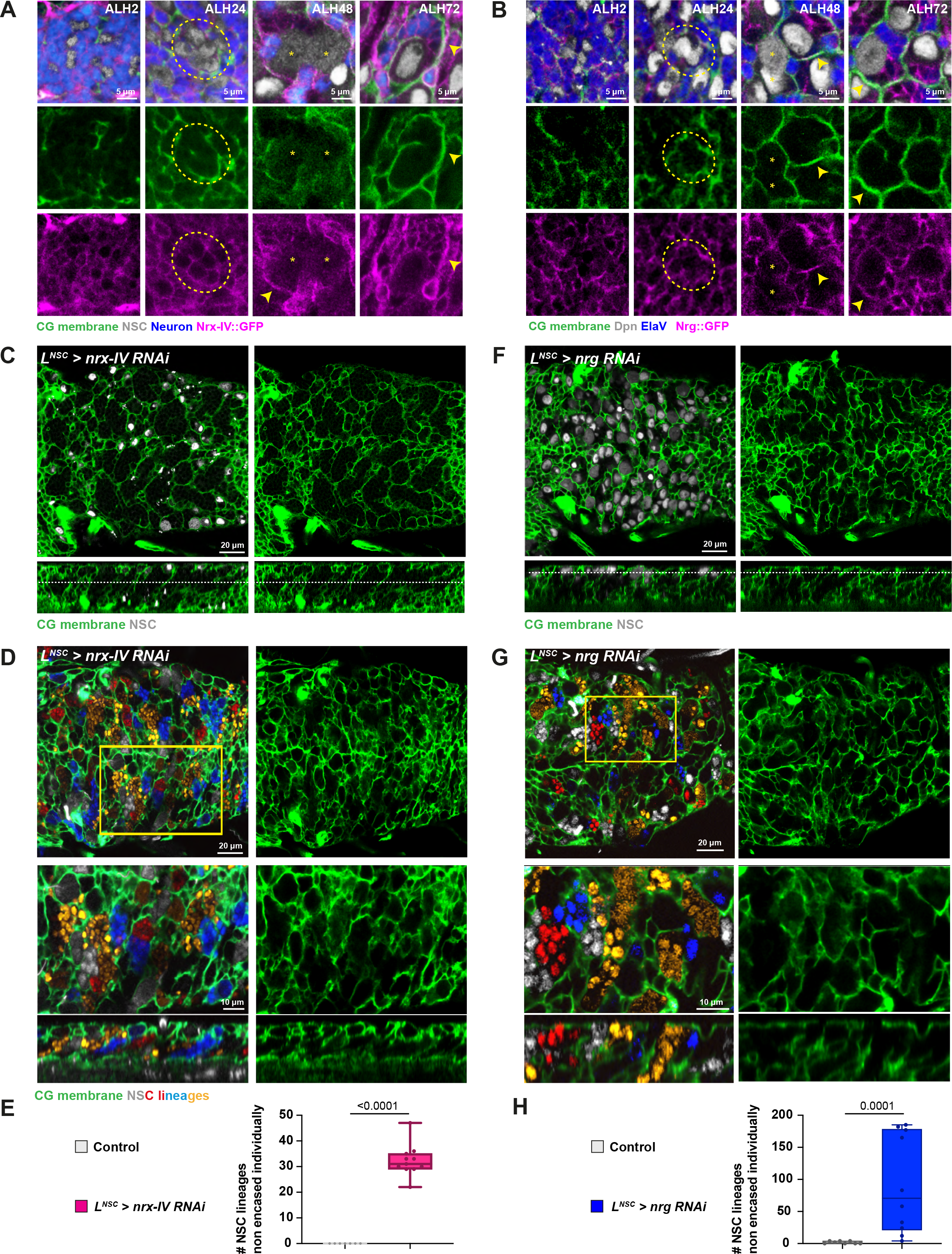
Neurexin-IV and Neuroglian are both required in NSC lineages for individual encasing. A) Representative confocal images of the expression of Neurexin-IV (Nrx-IV) at ALH0, ALH24, ALH48 and ALH72 at 25°C. Nrx-IV is monitored through a *Nrx-IV∷GFP* fusion (magenta), CG membrane is visualized by *Nrv2∷GFP* (green), NSCs are labelled with Dpn (grey), and neurons are labelled with ElaV (blue). B) Representative confocal images of the expression of Neuroglian (Nrg) at ALH0, ALH24, ALH48 and ALH72 at 25°C. Nrg is monitored through a *Nrg∷GFP* fusion (magenta), CG membrane is visualized by *Nrv2∷GFP* (green), NSCs are labelled with Dpn (grey), and neurons are labelled with ElaV (blue). C) Representative confocal picture (median and orthogonal views) of the thoracic VNC for a condition in which *nrx-IV* is knocked down by RNAi from ALH0 in NSC lineages (*L^NSC^ > nrx-IV RNAi*, driver line *Nrv2∷GFP, wor-GAL4; tub-GAL80^ts^*). Larvae are dissected after 68 h at 29°C. CG membrane is visualized by *Nrv2∷GFP* (green) and NSCs are labelled with Dpn (grey). The dashed white line on the orthogonal view indicates the plane chosen for the median view. D) Representative confocal picture of the thoracic VNC for a condition in which *nrx-IV* is knocked down by RNAi from ALH0 in NSC lineages marked with the multicolour lineage tracing Raeppli-NLS (blue, white, orange and red). Raeppli-NLS is induced at ALH0 using *hs-Flp*. CG membrane was visualized with *Nrv2∷GFP* (green). See Methods for timing, conditions and genetics of larval rearing. E) Quantification of the number of NSC lineages non-individually encased from D). Control (n = 7 VNCs) and *nrx-IV RNAi* (n = 11 VNCs). Data statistics: Mann-Whitney test. Results are presented as box and whisker plots. F) Representative confocal picture (median and orthogonal views) of the thoracic VNC for a condition in which *nrg* is knocked down by RNAi in NSC lineages (*L^NSC^ > nrg RNAi*, driver line *Nrv2∷GFP, wor-GAL4; tub-GAL80^ts^*). Larvae are dissected after 24 h at 18°C followed by 54 h at 29°C. CG membrane is visualized by *Nrv2∷GFP* (green) and NSCs are labelled with Dpn (grey). The dashed white line on the orthogonal view indicates the plane chosen for the median view. This phenotype is seen in 7/10 cases, 3/10 show a milder phenotype. G) Representative confocal pictures of the thoracic VNC for a condition in which *nrg* is knocked down by RNAi in NSC lineages marked with the multicolour lineage tracing Raeppli-NLS (blue, white, orange and red). Raeppli-NLS is induced at ALH0 using *hs-Flp*, and RNAi after 24 h at 18°C. Larvae are dissected 54 h after RNAi induction. CG membrane was visualized with *Nrv2∷GFP* (green). See Methods for timing, conditions and genetics of larval rearing. H) Quantification of the number of NSC lineages non-individually encased from G). Control (n = 8 VNCs) and *nrg RNAi* (n = 10 VNCs). Data statistics: Mann-Whitney test. Results are presented as box and whisker plots.

Strikingly, the enrichment at the interface between cells from the same NSC lineage was also present for other SJ components, namely Dlg1, ATPα and Cora, with the latter also exhibiting a staining along the CG interface (Supp. Fig. 4A). We were not able to assess Cont due to lack of working reagents. Taken together, these data suggest that multiple SJ components are expressed in NSC lineages, localizing between cells of the same lineages, as well as between lineages and the CG.

### Nrx-IV and Nrg are required in NSC lineages for individual encasing by CG

We then asked the importance of such expression in the individual encasing of NSC lineages by CG. We decided to knock down from ALH0 *nrx-IV*, *nrg*, *dlg1*, *cont* and *ATPα* in NSC lineages using specific RNAi lines. Larvae from *dlg1*, *cont* and *ATPα* knockdown died at early larval stages. From ATPα knockdown, few larvae still reached late larval stage, displaying restricted irregularities the CG network (Supp. Fig. 4B). In contrast, *nrx-IV* and *nrg* knockdowns, which successfully reduced the expression in NSC lineages of *Nrx-IV∷GFP* and *Nrg∷GFP*, respectively (Supp. Fig. 4C-D), both resulted in altered encasing of individual NSC lineages.

Under *nrx-IV* knockdown, CG pattern first appeared mostly normal when observed at the NSC level, with NSCs seemingly individually separated by CG membranes. However, we observed a striking, unusual pattern at the level of differentiating progeny, with much larger CG chambers harbouring a clear continuous membrane outline with no signal inside (Fig. 4C). A similar result was obtained when *nrx-IV* RNAi was expressed under the control of the neuronal driver ElaV-GAL4 (Supp Fig. 4E). In contrast, no effect was detected under *nrx-IV* knockdown in CG (Supp. Fig. 4E). Expressing the multicolour clonal marker Raeppli-NLS along with *nrx-IV* RNAi in NSC lineages (both induced at ALH0) revealed an extensive loss of individual encasing, with multiple NSC lineages not separated by CG membranes but rather clustered together in large chambers (Fig. 4D-E).

Driving *nrg* knockdown from larval hatching (ALH0) led to few larvae of the right genotype, in which CG displayed some restricted defects in individual encasing of NSC lineages. We thought these animals might have survived due to a weak phenotype, and decided to delay the RNAi knockdown, started after one day, and then maintained for 2 to 3 days. In this case, we obtained more surviving larvae which displayed strong alterations of NSC individual encasing by CG. At the stem cell level, NSC were indeed clustered together, seemingly touching each other (Fig. 4F). Going deeper at the level of the maturing progeny also revealed bigger zones devoided of CG membrane (see orthogonal view). Of note, the expressivity of the phenotype was variable (around 30% of the larvae display milder alterations, a population seen and representing the lower points in Fig. 4H). Driving *nrg* RNAi in neurons (ElaV-GAL4) in the same induction conditions also led to defects in the encasing of NSC lineages, albeit in a weaker fashion than with the NSC driver (Supp Fig. 4F). Finally, *nrg* knockdown in CG did not lead to observable CG alteration (Supp. Fig. 4F). Expressing Raeppli -NLS along with *nrg* RNAi in NSC lineages confirmed the loss of individual encasing, with multiple NSC lineages grouped together in large chambers, in a seemingly random fashion (Fig. 4G-H).

Altogether, our findings suggest that the expression of Nrx-IV and Nrg in NSC lineages are both required for their individual encasing by CG.

### A glia to NSC lineages adhesion through Nrx-IV and Wrapper is required for individual encasing

The dual role of Nrx-IV within and outside septate junctions is sustained by the existence of alternative splicing [46]. Nrx-IV can be produced as a SJ isoform (Nrx-IV^exon3^), and a neuronal isoform outside of SJ (Nrx-IV^exon4^). Nrx-IV role within the embryonic CNS is through its recruitment by and binding to its glial partner Wrapper, another member of the immunoglobulin family [45–47]. We thus sought to assess whether the role of Nrx-IV in NSC lineages encasing by CG was through, or independent of, Wrapper.

Previous studies had reported that regulatory sequences in the *wrapper* gene drive in the CG during larval stages [38,54]. Staining with an anti-Wrapper antibody confirmed Wrapper expression in CG, where it localizes in the membrane (Fig. 5A). In addition, a CRIMIC line [55], in which a CRISPR-directed intronic insertion allows *GAL4* expression under control of endogenous *wrapper* promoter, was used to drive fluorescent membrane (*UAS-mCD8∷GFP*) and nuclear (*UAS-His∷mRFP*) reporters. It produced the stereotypic CG pattern and co-stained with a glial fate marker, Repo (Supp. Fig. 5A). We then found that RNAi knockdown of *wrapper* in the CG reproduced the highly characteristic pattern of large chambers (Fig. 5B) found during Nrx-IV knockdown in NSC lineages (compare with Fig. 4C). Taken together, our results suggest that Nrx-IV in NSC lineages partners with Wrapper in the CG, outside of a septate junction function. Moreover, this interaction is essential to produce individual encasing of NSC lineages by CG, implying that in absence of such recognition, CG cannot properly recognize and individually sort NSC lineages.

**Figure 5.**
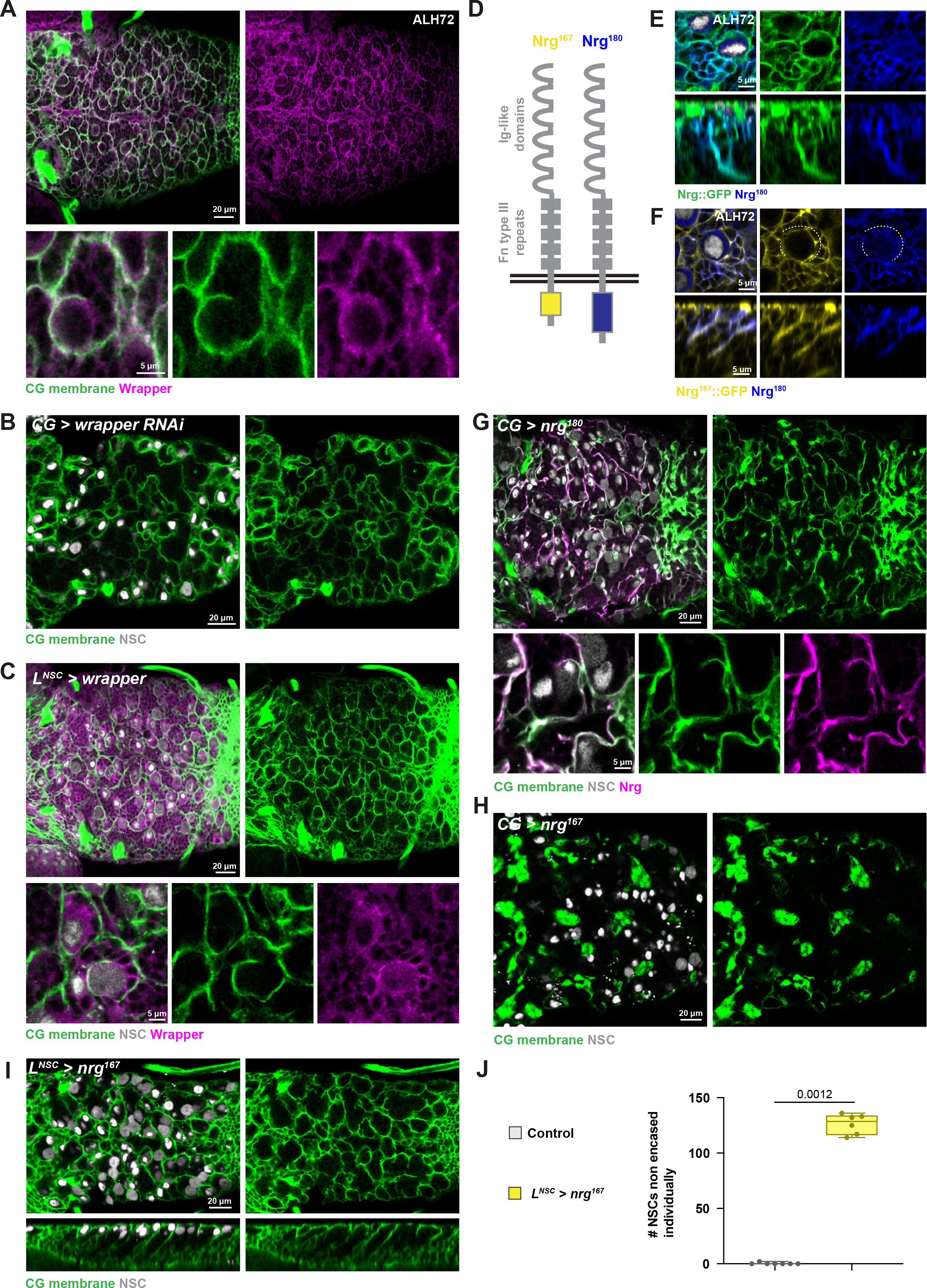
Individual encasing relies on balancing a CG to lineage interaction through Nrx-IV and Wrapper with an intra-lineage adhesion through Nrg. A) Representative confocal picture of the localisation of Wrapper in a thoracic VNC. Larvae were dissected at ALH72 at 25°C. CG membrane is visualized by *Nrv2∷GFP* (green) and Wrapper is detected by a specific antibody (magenta). B) Representative confocal picture of the thoracic VNC for a condition in which *wrapper* is knocked down by RNAi in the CG (*CG > wrapper RNAi*, driver line *Nrv2∷GFP, tub-GAL80^ts^; cyp4g15-GAL4*). Larvae are dissected after 68 h at 29°C from ALH0. CG membrane is visualized by *Nrv2∷GFP* (green) and NSCs are labelled with Dpn (grey). C) Representative confocal picture of the thoracic VNC for a condition in which *wrapper* is overexpressed in NSC lineages from ALH0 (*L^NSC^ > wrapper*, driver line *Nrv2∷GFP, wor-GAL4; tub-GAL80^ts^*). Larvae are dissected after 68 h at 29°C. CG membrane is visualized by *Nrv2∷GFP* (green) and NSCs are labelled with Dpn (grey). D) Schematic depicting the two isoforms for Nrg, Nrg^167^ and Nrg^180^. Only the intracellular C-terminal part differs. E) Representative confocal picture of the localisation of the Nrg^180^ isoform in a thoracic VNC, at ALH72 at 25°C. All Nrg isoforms are monitored through a *Nrg∷GFP* protein trap (green) and the Nrg^180^ isoform is detected with a specific antibody (BP104, blue). F) Representative confocal close-up picture of the respective localisations of the Nrg^167^ and Nrg^180^ isoform in a thoracic VNC, at ALH72 at 25°C. The Nrg^167^ isoform is visualized by a protein trap in the *nrg* gene leading to the preferential expression of this isoform (*Nrg^167^∷GFP*, yellow). The Nrg^180^ isoform is detected with a specific antibody (BP104, blue). The dashed white line highlights the perimeter of the NSC devoided of BP104 signal. G) Representative confocal picture of a thoracic VNC and close-up for Nrg^180^ overexpression in the CG from ALH0 (*CG > nrg^180^*, driver *Nrv2∷GFP, tub-GAL80^ts^; cyp4g15-GAL4*). Larvae are dissected after 68 h at 29°C. CG membrane is visualized by *Nrv2∷GFP* (green), NSCs are labelled with Dpn (grey) and Nrg^180^ is detected with a specific antibody (BP104, magenta). H) Representative confocal picture of a thoracic VNC for Nrg^167^ overexpression in the CG from ALH0 (*CG > nrg^167^*, driver *Nrv2∷GFP, tub-GAL80^ts^; cyp4g15-GAL4*). Larvae are dissected after 68 h at 29°C. CG membrane is visualized by *Nrv2∷GFP* (green) and NSCs are labelled with Dpn (grey). I) Representative confocal picture of a thoracic VNC for a condition in which *nrg^167^* is overexpressed from ALH0 in NSC lineages (*L^NSC^ > nrg^167^*, driver line *Nrv2∷GFP, wor-GAL4; tub-GAL80^ts^*). Larvae are dissected after 68 h at 29°C. CG membrane is visualized by *Nrv2∷GFP* (green) and NSCs are labelled with Dpn (grey). J) Quantification of the number of NSCs non-individually encased from I). Control (n = 7 VNCs) and *nrg^167^* (n = 6 VNCs). Data statistics: Mann-Whitney test. Results are presented as box and whisker plots.

We then wondered how the Nrx-IV to Wrapper interaction would fit in the hypothesis of a difference in adhesion between A_L_ and A_L-CG_. If CG to NSC lineage adhesion is indeed weaker than intra-lineage adhesion (Fig. 1H, panel II.2), shifting Nrx-IV binding to Wrapper from a glial to a NSC lineage pool (thus providing intra-lineage adhesion through Nrx-IV and Wrapper in addition to other existing interactions) should not affect the sorting between NSC lineages and CG, since A_L_ would still be superior to A_L-CG_. However, if Nrx-IV to Wrapper interaction is stronger than the sum of intra-lineage adhesions, then forcing it in the lineage would tend to flatten the difference in adhesion (A_L_ ≈ _AL-CG_), and thus lead to random gouping of NSC lineages together. We found that misexpressing *wrapper* in NSC lineages from larval hatching (ALH0), while successful, resulted in very little alteration of CG encasing of individual NSC lineages (Fig. 5C). These data plead in favour of a CG to NSC lineage adhesion through Nrx-IV and Wrapper being weaker than the sum of intra-lineage adhesions.

### Intra-lineage adhesion through Nrg drives individual encasing by CG

Similarily to Nrx-IV, the dual role of Nrg in and outside of septate junction bores from differential splicing [56]. Nrg indeed comes in two isoforms, with the same extracellular domain but different intracellular parts (Fig. 5D). While the short isoform, Nrg^167^, localizes in the SJ of epithelial tissues, the long isoform, Nrg^180^, is expressed in neurons of the central and peripheral nervous systems during development [50,57,59]. We first determined which isoform is expressed in NSC lineages during larval stage, taking advantage of an antibody (BP104) specifically recognizing the Nrg^180^ isoform [56]. Staining of *Nrg∷GFP* CNS (ALH72) with BP104 revealed that Nrg^180^ localises in the membranes of all cells from NSC lineages (Fig. 5E and Supp. Fig. 5B). In neurons, it was not only found in the cell body but also in the axonal bundle, a localisation reported previously [53]. In accordance with this result, *nrg* knockdown in NSC lineages completely depleted the BP104 signal (Supp. Fig. 5C). We then took advantage of a Nrg∷GFP fusion which has been shown in other tissues to preferentially target the Nrg^167^ isoform (called *Nrg^167^∷GFP*, [48,50]). *Nrg^167^∷GFP* also appeared enriched between cells of the same NSC lineage, where it co-localised with BP104 staining, except on the NSC perimeter, devoided of BP104 (Fig. 5F and Supp. Fig. 5D; see dashed white line for lack of BP104). In contrast, only Nrg^167^ is detected in septate junctions. Taken together these data suggest that the two isoforms of Nrg are expressed in NSC lineages.

While Nrg can also bind in an heterophilic manner, mostly homophilic interactions (between same or different isoforms) have been reported. We thus wondered whether an Nrg to Nrg interaction within the NSC lineages could fulfill the role of an intra-lineage adhesion stronger than an CG to NSC adhesion. First, *nrg* knockdown in CG, the only cell population in contact with NSC lineages (besides clonally-related cells) did not recapitulate *nrg* knockdown in NSC lineages (compare Fig. 4F with Supp. Fig. 4F), suggesting that homophilic Nrg interactions do not exist between CG and NSC lineages to maintain individual encasing.

We then assessed the relevance of intra-lineage Nrg interactions in setting up a differential in adhesion (Fig. 1H, panel II.2). If such adhesion is stronger than the CG to NSC lineage interaction, then expressing Nrg in CG would force CG to interact with each other (A_L-CG_ through Nrg would be less favoured as a higher dose of Nrg would be present in CG due to overexpression). Strikingly, misexpressing Nrg^180^ in CG from larval hatching (ALH0) resulted in altered CG morphology and loss of individual encasing of NSC lineages (Fig. 5G). CG membranes displayed local accumulation as well as unusual curvature, and NSCs were not separated from each other by CG anymore but were rather found grouped close to each other. Overexpression of Nrg^167^ in CG (from ALH0) produced an even more dramatic phenotype, with localised, compact globules of CG membranes and the complete lack of individual encasing of NSC lineages (Fig. 5H). Interestingly, misexpression of a Nrg^GPI^ construct in which the transmembrane and cytoplasmic domains are replaced by a GPI anchor signal [58] also resulted in aggregated CG and clustered NSC lineages (Supp. Fig. 5E). This shows that intracellular signalling through the divergent C-terminal domain is not required for this sorting of CG and NSC lineages, but rather that adhesion through the extracellular part mediates this effect.

Altogether, these results demonstrate that forcing CG to CG adhesion through Nrg homophilic interactions is sufficient to segregate them from the whole population of NSC lineages, between which weaker interactions exist. This further suggests that Nrg homophilic adhesions between cells of the same NSC lineage are responsible for keeping these cells together and excluding CG.

If Nrg interactions are responsible for providing binding between cells of the same NSC lineage, including the stem cell, one consequence is that NSC could bind to each other. That would result in several NSCs encased in the same CG chamber something we do not witness in normal conditions. Interestingly, Nrg seems to be expressed in NSC after their encasing (see Fig. 4B), in contrast to Nrx-IV (Fig. 4A). This would fit with the idea that early on, when NSC are not encased yet and separated from other NSCs by the CG, A_NSC-NSC_ is kept low. As such, a precocious expression of Nrg in NSCs would be predicted to lead to their grouping (and further the grouping of their lineages) in a CG chamber. Strikingly, expressing Nrg^167^ (which gave the strongest phenotype in CG, see Fig. 5G-H) from ALH0 resulted in multiple CG chambers containing several NSCs (Fig. 5I-J). A similar result was obtained when expressing Nrg^GPI^ with the same timing (Supp. Fig. 5F), implying that the adhesive role of Nrg is responsible for such effect. This is in contrast to the lack of effect of misexpressing Wrapper in NSC lineages (also from ALH0), showing that not all adhesion complexes can lead to A_NSC-NSC_ high enough to group NSC together. These results suggest that a proper timing in establishing intra-lineage adhesion is instrumental in ensuring the individual encasing of NSC lineages by CG.

### Nrx-IV and Nrg adhesions are required in NSC lineages for correct axonal path during development

So far, our data show that Nrx-IV- and Nrg-mediated adhesions in NSC lineages, while likely fulfilling different roles in this process, are both important to set up the individual encasing of NSC lineages by CG. We sought to assess the functional relevance of such adhesions, and ultimately of individual encasing, for the cells of NSC lineages. We first turned our eyes to the NSC themselves. As NSC core function is dividing to produce differentiated progeny, we assessed NSC proliferation under *nrg* and *nrx-IV* knockdown in NSC lineages, using phospho-histone 3 (PH3) to mark mitotic DNA. We found that both mitotic indexes and phase distribution in mitosis were similar between these conditions and control (Supp. Fig. 6A-B). Of note, *shg* knockdown in NSC lineages also did not lead to detectable changes in the mitotic profile (Supp. Fig. 6C). These results show that Nrg and Nrx-IV adhesion in the niche are not critical for the rate of NSC proliferation

Improper encasing of NSC lineages during development implies that newborn neuronal lineages are not physically constrained anymore nor neatly packed. Rather they are expanding more freely and are found mingled with other ones. At this stage, immature secondary neurons start sending axons to establish synaptic connections with proper partners in the neuropile, with axons from the same lineage grouped as one or two tight bundles and following the same path [27,53] (Supp. Fig. 1A and Fig. 6A). This axonal fascicle, also encased by the CG membrane, shows a well-defined tract for each lineage, with stereotyped entry in and path within the neuropile.

**Figure 6.**
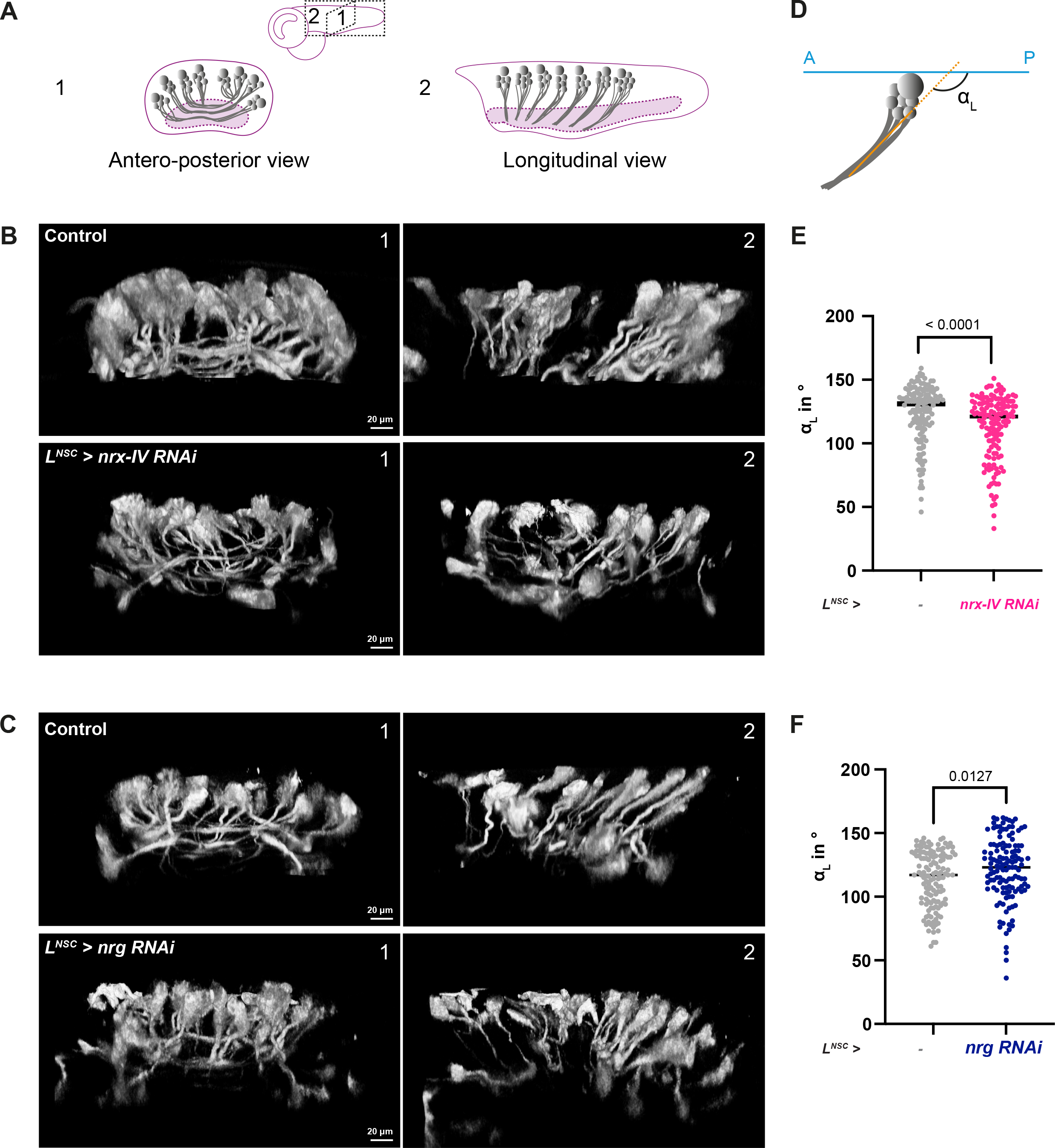
Loss of Nrx-IV and Nrg adhesions in NSC lineages during development induces axonal misprojection from newborn neurons. A) Schematic of the axonal projections coming from secondary, newborn neurons generated by NSCs during larval neurogenesis. Only the VNC region is depicted. 1, antero-posterior view. 2, longitudinal view. B) 3D reconstruction of a group of NSC lineages visualized with a membrane marker (mTFP1-CAAX) in antero-posterior (1) and longitudinal (2) views for a control condition and for *nrx-IV* knockdown in NSC lineages (*L^NSC^ > nrx-IV RNAi*). Clonal labelling was obtained through the induction of Raeppli-CAAX in NSC lineages at ALH0. See Methods for timing, conditions and genetics of larval rearing. C) 3D reconstruction of a group of NSC lineages visualized with a membrane marker (mTFP1-CAAX) in antero-posterior (1) and longitudinal (2) views for a control condition and *nrg* knockdown in the NSC lineages (*L^NSC^ > nrg RNAi*). Clonal labelling was obtained through the induction of Raeppli-CAAX in NSC lineages at ALH0. See Methods for timing, conditions and genetics of larval rearing. D) Schematic of the angle between the main axonal tract projecting from secondary newborn neuron and the antero-posterior axis. E) Quantification of the angle α_L_ depicted in D) in VNCs for control and *nrx-IV* knockdown in NSC lineages, in the same conditions shown in B). Control (n = 149 axonal projections from 7 VNCs) and *nrx-IV* RNAi (n = 143 axonal projections from 8 VNCs). Data statistics: Mann-Whitney test. Results are presented as individual values, the line represents the median. F) Quantification of the angle α_L_ depicted in D) in VNCs for control and *nrg* knockdown in NSC lineages, in the same conditions shown in C). Control (n = 144 axonal projections from 7 VNCs) and *nrg* RNAi (n = 129 axonal projections from 6 VNCs). Data statistics: Mann-Whitney test. Results are presented as individual values, the line represents the median.

We thus wondered whether disruption of niche adhesion and loss of lineage organization could translate into an altered pattern of axonal projections. To assess this possibility, we marked NSC lineages in a multicolour clonal fashion, this time using a membrane version of Raeppli (CAAX tag) [34] to label both the cell body as well as the extending axons (Supp. Fig. 1A and Fig. 6B-C). Both for *nrx-IV* and *nrg* knockdowns, we first found that the organization in bundles of axons from neurons of the same lineage appeared preserved, and that most of them still found their way to the neuropile. We however noticed a less regular pattern in their path to the neuropile, drifting from the classic boat shape seen from the antero-posterior view (Fig. 6A-C, view 1) and appearing less aligned in a longitudinal view (Fig. 6A-C, view 2). We then calculated the angle of axonal extension to the antero-posterior axis of the VNC (Fig. 6D). We found that, compared to a control condition at the same stage (slightly earlier for *nrg* RNAI), the angles were less stereotyped in overall, with a broader distribution, and slightly shifted, being either more closed (*nrx-IV* RNAi, Fig. 6E) or more open (*nrg* RNAi, Fig. 6F). In addition, we also detected the rare occurrence of axonal bundles not targeting the neuropile, but rather going to the edge of the organ, without establishing synaptic connections (Supp. Fig. 6D). Altogether, these data show that Nrx-IV and Nrg adhesions in the NSC niche both influence the extension of axonal tracts from newborn neurons.

### The function of Nrx-IV and Nrg adhesions in NSC lineages during development influences adult locomotor behaviour

The tracts of the axonal projections established by secondary neurons during larval development are mostly kept during metamorphosis, being extended and complexified rather than fully remodeled [26,27]. Indeed, despite the overall change in CNS morphology over time, the relative positions and pattern of tracts from different lineages are maintained and recognizable [53]. As such, the correct establishment of axonal projections during development is meant to be critical for the function of mature neurons in the adult CNS. In this light, we decided to determine whether the loss of Nrx-IV and Nrg adhesions in NSC lineages during development could impair neuronal function later in the adult.

One way to assess neuronal function is to probe its functional output on adult behaviour. As our analysis of CG and axonal tract phenotypes have focused on the VNC, in which motor neurons are produced, we focused on motor parameters in the adult. To do so, we took advantage of an ethoscope-based tracking system [60] to record locomotion metrics. This high-throughput platform relies on video acquisition to record positional data in real-time for multiple flies, individually placed in a cylindrical tube of given length and volume. Several behavioural parameters can be extracted by calculating the position of the fly overtime, including average locomotion speed (velocity), amount of sleep (defined as the cumulative time during which a fly stay still for at least five minutes, [61]) and circadian activity. Statistics on several flies finally allow to draw an average behaviour for the population.

We recorded locomotion metrics for *shg*, *nrx-IV* and *nrg* knockdowns in NSC lineages, as well as for a control line. We did it in two conditions. First, RNAi expression was only allowed during larval phase, and prevented from mid-pupal stage (see Methods, “induced” condition). In this case, comparing the behaviour of the different knockdowns to the control line reveals the importance of each adhesion on locomotion parameters. Second, gene knockdowns were never activated (same genetic background, but RNAi always off; see Methods, “Non Induced” condition), a condition meant to serve as a control for the effect of genetic background on locomotion parameters. We did not find comparing induced and non-induced relevant, as these two conditions relies on very different regimens of fly husbandry.

We first look at the overall pattern of activity through a circadian cycle of light and dark periods. In laboratory conditions, *Drosophila* indeed displays a characteristic rest (sleep)/activity pattern where they become highly active in anticipation of the transitions between light and dark periods. Rest/sleep happens mostly in the middle of light and dark periods. We found that this circadian pattern of activity was kept in the different lines in induced condition, with two main peaks of activity (morning and evening, Supp. Fig. 7A). We noticed slightly higher anticipation for the evening peak in the case of *nrx-IV* and *nrg* knockdowns, as well as wider peaks for *nrg* knockdown. Sleep metrics then revealed a stunning change in the behaviour of *nrg* RNAi and *nrx-IV* RNAi flies, while *shg* RNAi appeared very similar to control (Fig. 7A, induced). While control and *shg* RNAi flies were spending 70% and 68% of their time sleeping, respectively, *nrg* RNAi flies spent only 27% of their time in average, a dramatic reduction (Fig. 7B, induced). They appeared hyperactive throughout both light and dark periods (Supp. Fig. 7B), with an especially important shift during the time between activity peaks. *nrx-IV* RNAi flies also spend significantly less time sleeping, which was decreased to only 58 % of their time (Fig. 7B, induced). In contrast, *shg* RNAi, *nrx-IV* RNAI, *nrg* RNAi and control all behaved in a similar fashion in the non-induced condition (Fig. 7A-B; ctrl = 61%; *shg* = 53%; *nrx-IV* = 54%; *nrg* = 58% of time sleeping).

**Figure 7.**
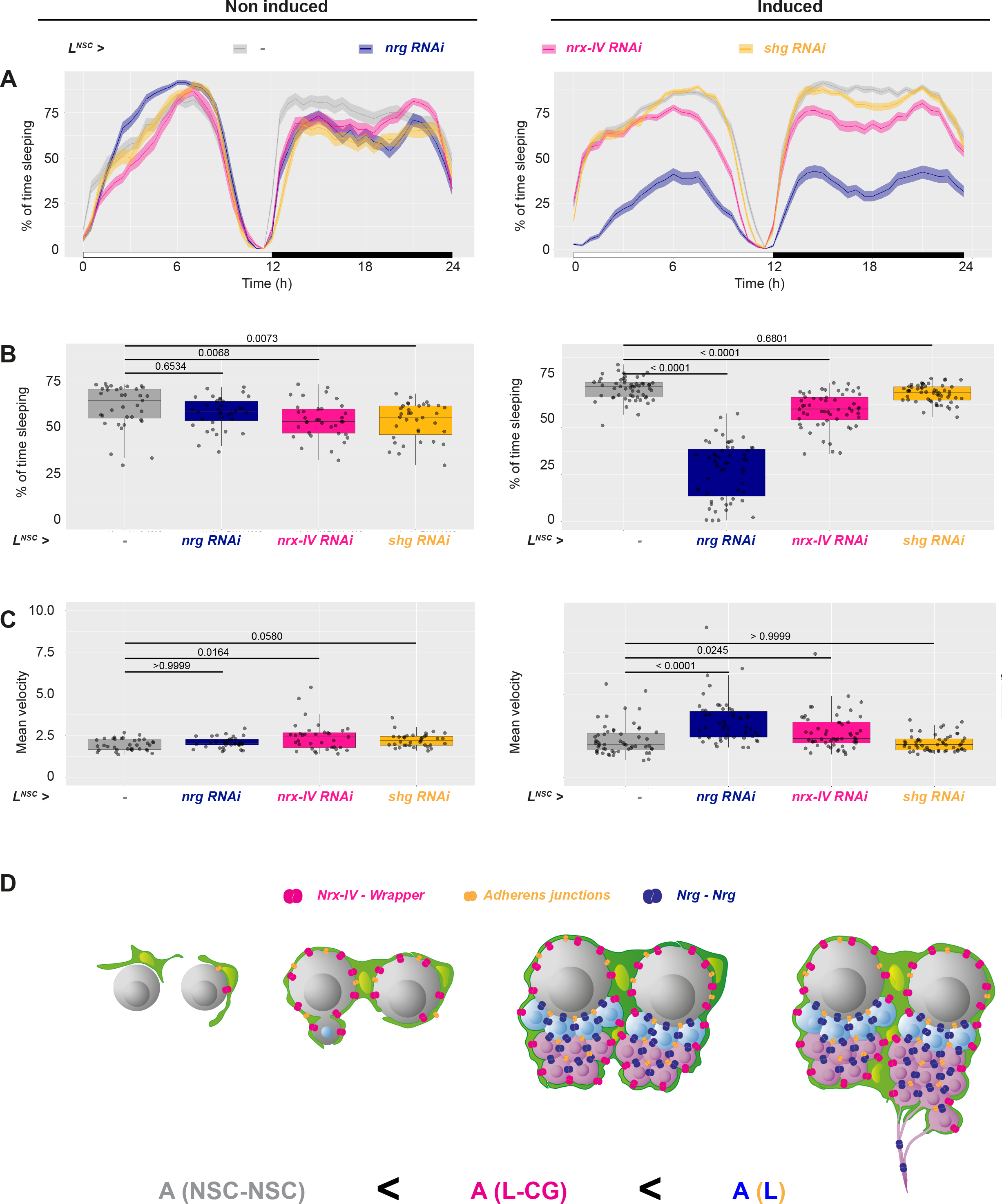
Loss of Nrx-IV and Nrg adhesions in NSC lineages during development results in locomotor hyperactivity in the resulting adults. A) Plot representing the percentage of global time sleeping (% asleep, ratio between total sleep time and total time), in non-induced (flies always kept at 18°C before the recordings) and induced (flies shifted to 29°C from early larval stage to mid-pupal stage) conditions. Control, (*wor-GAL4, tub-Gal80^ts^ x w^1118^*), n = 35 non-induced adult males and n = 55 induced adult males. *shg* RNAi (*wor-GAL4, tub-Gal80^ts^ x shg RNAi^VDRC27082^*), n = 35 non-induced adult males and n = 55 induced adult males. *nrx-IV* RNAi (*wor-GAL4, tub-Gal80^ts^ x nrx-IV RNAI^BL32424^*), n = 35 non-induced adult males and n = 55 induced adult males. *nrg* RNAi (*wor-GAL4, tub-Gal80^ts^ x nrg RNAI^BL37496^*), n = 35 non-induced adult males and n = 55 induced adult males. B) Fraction (%) of the time sleeping across Light/Dark cycles (measured as the fraction of time sleep within 30 min intervals) in non-induced (flies always kept at 18°C before the recordings) and induced (flies shifted to 29°C from early larval stage to mid-pupal stage) conditions. Control, (*wor-GAL4, tub-Gal80^ts^ x w^1118^*), n = 35 non-induced adult males and n = 55 induced adult males. *shg* RNAi (*wor-GAL4, tub-Gal80^ts^ x shg RNAi^VDRC27082^*), n = 35 non-induced adult males and n = 55 induced adult males. *nrx-IV* RNAi (*wor-GAL4, tub-Gal80^ts^ x nrx-IV RNAI^BL32424^*), n = 35 non-induced adult males and n = 55 induced adult males. *nrg* RNAi (*wor-GAL4, tub-Gal80^ts^ x nrg RNAI^BL37496^*), n = 35 non-induced adult males and n = 55 induced adult males. Data statistics: one-way ANOVA with a Kruskal–Wallis multiple comparison test. Results are presented as box and whisker plots. C) Mean velocity (in relative units) across Light/Dark cycles in non-induced (flies always kept at 18°C before the recordings) and induced (flies shifted to 29°C from early larval stage to mid-pupal stage) conditions. Control, (*wor-GAL4, tub-Gal80^ts^ x w^1118^*), n = 35 non-induced adult males and n = 55 induced adult males. *shg* RNAi (*wor-GAL4*, *tub-Gal80^ts^ x shg RNAi^VDRC27082^*), n = 35 non-induced adult males and n = 55 induced adult males. *nrx-IV* RNAi (*wor-GAL4, tub-Gal80^ts^ x nrx-IV RNAI^BL32424^*), n = 35 non-induced adult males and n = 55 induced adult males. *nrg* RNAi (*wor-GAL4, tub-Gal80^ts^ x nrg RNAI^BL37496^*), n = 35 non-induced adult males and n = 55 induced adult males. Data statistics: one-way ANOVA with a Kruskal–Wallis multiple comparison test. Results are presented as box and whisker plots. D) Schematic depicting the timing and localisation of different adhesion complexes within the NSC niche which are required for the individual encapsulation of NSC lineages by CG. While Nrx-IX starts to be expressed in NSCs before encapsulation, Nrg only appears afterwards, a timing preventing the clustering of NSCs, and hence later on, NSC lineages, within one chamber. Nrg binds to itself in the NSC lineages (dark blue complexes), while Nrx-IV binds to Wrapper expressed in the CG (pink complexes). Adherens junctions (orange complexes) are also present between the cells of the same NSC lineage, where they are mostly dispensible for individual encasing, while potentially providing robustness.

We wondered whether this locomotor hyperactivity was only visible as the time flies spent moving, or also in the way they were moving. We then determined the speed of locomotion for the different lines. In induced conditions, we found that the mean velocity throughout the cycle was increased in *nrg* (3.5) and *nrx-IV* (2.8) but not *shg* (2.1) knockdowns compared to control condition (2.3). In non-induced conditions, all lines exhibited similar values of velocity (ctrl = 2.0; *shg* = 2.2; *nrx-IV* = 2.4; *nrg* = 2.1). Taken together, these results show that the functions of Nrg and and Nrx-IV, two adhesion molecules required for the individual encasing of NSC lineages, are also necessary during development for proper motor activity in the adult.

## DISCUSSION

The NSC niche harbours an elaborate architecture surrounding the stem cells and their growing, differentiating neuronal lineages. However, its mechanisms of formation and its role on NSCs and newborn progeny remain poorly understood. Here we investigate the formation of glial niches around individual NSC lineages in the *Drosophila* developing CNS. Individual encasing occurs around the NSC itself, before neuronal production, providing a first mechanism for implementing lineage encasing. However, such timing is not the only strategy to ensure the formation and maintenance of individual encasing around the entire lineage. We actually uncovered that CG are able to distinguish between and sort themselves from individual NSC lineages through differences in adhesion complexes, what provides a belt and braces mechanism to ensure lineage encasing regardless of timing. Several components of both adherens and occluding junctions are indeed expressed in NSC lineages. While adherens junctions appear mostly dispensible for lineage encasing, two SJ components, Nrx-IV and Nrg, are required for this structure, however outside of their junctional roles. Nrx-IV binds to Wrapper present on the CG, and Nrg, expressed after neuronal production starts, perfoms homophilic interactions with itself to bind cells from one lineage together. This Nrg-based intra-lineage adhesion is instrumental in sorting NSC lineage and CG after neuronal production, providing a stronger adhesion compared to the Nrx-IV to Wrapper interaction. Finally, we found removing Nrx-IV and Nrg during larval stage leads to behavioural defects in adult, producing hyperactive flies. Altogether, our findings show that a timely difference in adhesion between NSC/NSC lineages and niche cells defines the structure of the niche during development and influences adult behaviour (Fig. 7D).

Both adherens and occluding junctions are associated with and as such mostly have been mostly described in epithelia and epithelial-like tissues. Here the fact that core components of adherens (E-cadherin, β-catenin) and occluding (Nrx-IX, Nrg, Dlg1, ATPa, Cora) junctions localise in stem cell and maturing progeny raises questions about their regulation and role in such cell types. While adherens junction appears to be specifically set up in NSC lineages, we did not find them functionally relevant for individual encasing by CG, NSC proliferation and motor behaviour in adult. Previous studies had reported that E-cadherin disruption was leading to defects in CG architecture and altering NSC proliferation. However, both studies used a dominant-negative form of E-cadherin [22,41]. Here, we could not recapitulate such consequences using an efficient RNAi knockdown nor a null allele of E-cadherin (Fig. 3C-D and Supp. Fig. 3C, G). It is possible that some of the effects observed previously are neomorphic and triggered by the activation of other pathways. Another possibility is the fact that knockdown, but not competition by a dominant-negative, could lead to compensation (such as an increase in N-cadherin), masking the role of E-cadherin and adherens junction. Nevertheless, while we believe adherens junctions are not instrumental in establishing CG architecture around NSC lineages, it might be used as a strengthening, safety mechanism, available when other processes fail. In this light, it would be relevant to determine whether *shg* knockdown potentializes *nrg* knockdown. Other roles in NSC lineages, possibly subtle, also await to be uncovered.

We propose that a balance between a strong Nrg-based adhesion within the NSC lineages and a weaker, Nrx-IV-based interaction between CG and NSC lineages (A_L-CG_ < A_L_) builds the stereotyped, individual encasing of NSC lineages. There is no direct mesure of the strength of adhesion between Nrx-IV and Wrapper compared to Nrg with itself. However, the fact that misexpressing Nrg in CG creates CG aggregates and alters individual encapsulation indicates that Nrg can surpass the endogenous Nrx-IV to Wrapper interaction. In this line, misexpressing Wrapper in the NSC lineages does not alter encasing, suggesting that increasing A_L_ compared to A_L-CG_ does not change the directionality of the difference, and as such that this difference is already there. Similarily, E-cadherin misexpression in the CG has no impact on lineage encasing, implying that the sum of the adhesions present in the NSC lineages outweights the presence of *de novo* adherens junctions between lineages and CG.

Nrx-IV interaction with Wrapper could be important to provide a scaffold onto which anchoring the glial membrane on the available surface of all lineage cells. When this scaffold is weakened, CG randomly infiltrate in between NSC lineages still tightly bound by Nrg interaction, leading to the creation of CG chambers of variable size (Fig. 4D-E). Such chambers appear neat, with a clear outline around grouped lineages and the absence of CG membrane signal within. Such striking, unmistakable phenotype, which we never observed previously, suggests that upon alteration of Nrx-IV and Wrapper interaction, CG still recognize NSC lineages as “wholes”, but cannot implement their individual encapsulation.

We propose that Nrg interact with itself in NSC lineages. We found that the Nrg^180^ isoform was expressed in newborn secondary neuron, in accordance with its known neuronal association in other life stages. However, we propose that Nrg^167^, traditionally associated with junctional localisation, is also present. This is based on the use of a Nrg∷GFP fusion shown to preferentially target the Nrg^167^ isoform. We also noticed that driving *nrg* RNAi in NSC lineages, under the same condition, completely abolished the signal from Nrg^180^ (BP104, Supp. Fig. 5C), but not from the *Nrg∷GFP* fusion (Supp. Fig. 4D. Whether Nrg^167^ interacts with Nrg^180^ for NSC encasing, or has another junctional role, remains to be demonstrated. Of note, an elegant study has shown that neuronal Nrg^180^ can bind to epidermal Nrg^167^ to prevent homologous Nrg^180^-mediated dendrite bundling and effectively promote enclosure of single-neuron dendrites by the epidermis [50], showing that isoform interactions exist to mediate cell-cell adhesion.

The temporal regulation of Nrg expression appears crucial. We found it is not present in NSCs before encapsulation (Fig. 4B), whereas its precocious expression results in NSC lineages grouped together (Fig. 5I-J and Supp. Fig. 5F). What triggers this timely change, and especially its link with NSC reactivating, is an intriguing question. Indeed, what first recruits CG membrane to NSC, before creating and clustering a lineage, remain to be identified.

The phenotypic expressivity of *nrg* knockdown in NSC lineages is variable, with a minority showing restricted defects, an observation we do not explain. Nevertheless, in most larvae, we find that *nrg* knockdown in NSC lineages result in multiple CG chambers with several NSC lineages grouped together, and in half of the case with only very few individual encasings left (Fig. 4H). In these conditions, cells from a same lineage still appear to be mostly kept together, as a group. If Nrg binds cells from the same lineage together, a potential outcome of its loss of function could have been for these cells to end up individually encapsulated, something we do not see. However, the existence of other adhesion complexes (such as E-cadherin) still providing cohesion might be enough to prevent the case where A_L_ < A_L-CG_ even under *nrg* knockdown. Ultimately, the total sum of adhesions for each cell pair decides of the directionality of the difference. We also do not know the adhesion strength between CG cells. If it is higher than the sum of remaining adhesions in NSC lineages after *nrg* knockdown, it could explain the fact that CG do not extend to separate NSC lineages.

The loss of Nrx-IV and Nrg adhesions both result in altered axonal tracts coming from the newborn neurons. Although it could be a consequence of some autonomous properties of these molecules on neuronal/axonal features, previous studies had already linked a change in CG structure or function to misshaped axonal tracts in the larval CNS, both in the larval optic lobe [17] or in the central brain, where genetic ablation of CG results in abnormal axonal trajectories and fasciculation [28].

We further linked the loss of Nrx-IV and Nrg adhesions during development to changes in the locomotor behaviour of the resulting adults, which appeared hyperactive. While Nrg could be involved in other ways than through building the niche structure, due its homotypic interactions between neurons and its strong axonal localization, Nrx-IV and Wrapper interaction makes a stronger case for bridging niche architecture with adult behaviour. First, Nrx-IV to Wrapper interaction is between lineages and CG, rather than between neurons. Moreover, in *nrx-IV* knockdown in NSC lineages and *wrapper* knockdown in CG, CG chambers are still present, neatly delineated around multiple NSC lineages, depicting the loss of the individuality of encasing rather than a comprehensive alteration of CG structure. This pleads for an impact of a targeted, specific remodelling of niche architecture during development on adult behaviour.

Hyperactivity (increased locomotor activity and reduced sleep) in *Drosophila* has been found in diverse models of neuro/developmental and neurological disorders, including the Fragile X syndrome [62], Attention-deficit/hyperactivity disorder (ADHD) [63] and Shwachman–Diamond syndrome [64]. In all these cases, changes in locomotion were correlated with synaptic and axonal abnormalities. Hyperactivity also appears during starvation, under hormonal control [65,66]. In our case, a reasonable explanation would be that axonal misprojection following loss of adhesion in the neurogenic niche during development translates to dysfunctional motor neurons in the adult.

Here, we propose a mechanism in which the temporal and spatial localization of different adhesion complexes results in the formation of a stereotypic niche organizing individual NSC lineages and their progeny. Their function is also important for axonal projection of newborn neurons, and locomotor behaviour in the adult, thus linking niche adhesive properties and developmental neurogenesis to adult health. All these complexes are heavily conserved in mammals, warranting the question of their non-junctional role in a developing CNS.

## METHODS

### Methods

#### Fly lines and husbandry

*Drosophila melanogaster* lines were raised on standard cornmeal food at 25°C. Lines used in this study are listed in the table below:

| Strains | Source | Stock number/Reference |
| --- | --- | --- |
| *w<sup>1118</sup>* | BDSC | 5905 |
| *Nervana2::GFP (Nrv2::GFP)* | BDSC | 6828 |
| *Shg::GFP* | Yohanns Bellaïche lab |  |
| *Nrx-IV::GFP* | Christian Klämbt lab | *Nrx<sup>454</sup>*, [67] |
| *Nrg::GFP* | Kyoto | 110658 |
| *Nrg<sup>167</sup>::GFP* | BDSC | 6844 |
| *Dlg1::GFP* | BDSC | 50859 |
| *ATPalpha::GFP (CPTI)* | Kyoto DGGR | 115323 |
| *tubulin-GAL80<sup>thermosensitive(ts)</sup>* | BDSC | 65406 |
| *yw, hs-FLP* | Andrea Brand lab |  |
| *FRT G13* | BDSC | 1956 |
| *CoinFLP* | BDSC | 58750 |
| *cyp4g15-GAL4* | BDSC | 39103 |
| *cyp4g15-FRT-STOP-FRT-GAL4* | This study |  |
| *cyp4g15-FRT-STOP-FRT-LexA* | Spéder lab | [25] |
| *cyp4g15-FLP* | Spéder lab | [25] |
| *cyp4g15-mtd::Tomato* | Spéder lab | [25] |
| *wor-Gal4* | BDSC (Doe lab insertions) | 56553 & 56554 |
| *PntP1-GAL4* | Jan lab | [68] |
| *CRIMIC wrapper* | BDSC | 93483 |
| *cyp4g15-QF2* | Spéder lab |  |
| *tub-QS* | BDSC | 52112 |
| *TUG G13 MARCM line*<br>*y,w, hs-FLP; FRTG13, tubP-GAL80[LL2]/*<br>*(CyO, act-GFP[JMR1]); tubP-GAL4[LL7],*<br>*UAS-mCD8-GFP[LL6]/TM6B* | Bruno Bello |  |
| *tubP-GAL80[LL2]* | BDSC | 5140 |
| *UAS-reaper* | Andrea Brand lab |  |
| *UAS-mCD8::GFP* | BDSC | 5130 |
| *UAS-mCD8::RFP* | BDSC | 27399 |
| *UAS-Raeppli CAAX 43E* | Generated from BDSC 55082 | [25] |
| *UAS-Raeppli NLS 53D* | Generated from BDSC 55087 | [25] |
| *UAS-Raeppli NLS 89A* | Generated from BDSC 55088 | [25] |
| *UAS-prospéro RNAi* | Andrea Brand lab |  |
| *UAS-brat RNAi* | Andrea Brand lab |  |
| *UAS-aPKc* |  | [69] |
| *UAS-shg RNAi* | BDSC | 32904 |
|  | VDRC | 27082 |
|  | VDRC | 103962 |
| *UAS-arm RNAi* | VDRC | 27227 |
| *UAS-nrx IV RNAi* | BDSC | 32424 |
| *UAS-nrg RNAi* | BDSC | 37496 |
| *UAS-wrapper RNAi* | BDSC | 29561 |
| *UAS-ATPα RNAi* | BDSC | 28073 |
| *UAS-nrg<sup>180</sup>* | BDSC | 24169 |
| *UAS-nrg<sup>167</sup>* | BDSC | 24172 |
| *UAS-nrg<sup>GPI</sup>* | BDSC | 24168 |
| *UAS-wrapper* | BDSC | 78535 |
| *QUAS-PTEN* | This study |  |
| *shg<sup>null</sup> (shg<sup>R64a</sup>)* | Yohanns Bellaïche lab |  |

#### Larval culture and staging

Embryos were collected within 2-4 hours window on grape juice-agar plates and kept at 25°C for 20-24 hours. Freshly hatched larvae were collected within a 1 hour time window (defined as 0 hours after larval hatching, ALH0), transferred to fresh yeast paste on a standard cornmeal food plate and staged to late first instar (ALH24), late second instar (ALH48), mid third instar (ALH72) and late third instar (ALH96).

For growth on quinic acid, food plates were prepared by mixing 250 mg/ml stock solution of quinin acid (dissolved in sterile water) into melted food at 50°C, for a final concentration of 20 mg/ml of quinic acid.

#### DNA cloning and *Drosophila* transgenics

A portion of the cyp4g15 enhancer (GMR55B12, Flybase ID FBsf0000165617), which drives in the cortex glia and (some) astrocyte-like glia, was amplified from genomic DNA extracted from cyp4g15-GAL4 adult flies, with a minimal Drosophila synthetic core promoter [DSCP] [70] fused in C-terminal. For creating cyp4g15-FRT-STOP-FRT-GAL4, a FRT STOP cassette was amplified from an UAS-FRT.STOP-Bxb1 plasmid (gift from MK. Mazouni) and the GAL4 sequence was amplified from the entry vector pENTR L2-GAL4∷p65-L5 (gift from M. Landgraf). The two amplicons were joined together by overlapping PCRs. This FRT-STOP-FRT-GAL4 amplicon together with the cyp4g15^DSCP^ enhancer were inserted in the destination vector pDESThaw sv40 using Multisite gateway system [71] to generate a cyp4g15^DSCP^-FRT-STOP-FRT-GAL4 construct. The construct was integrated in the fly genome at an attP2 or attP40 docking sites through PhiC31 integrase-mediated transgenesis (BestGene). Several independent transgenic lines were generated and tested, and one was kept for each docking site.

For creating *QUAS-PTEN*, the *PTEN* coding sequence was amplified from genomic DNA extracted from *UAS-PTEN* [72] adult flies, as described in [73]. This amplicon together with the QUAS sequence (pENTRY L1-QUAS-R5, gift from S.Stowers) were joined using the Multisite gateway system [71] in the destination vector pDESThaw sv40 gift from S. Stowers). The construct was integrated in the fly genome at an attP40 docking site through PhiC31 integrase-mediated transgenesis (BestGene). Several independent transgenic lines were generated and tested, and one was kept (*QUAS-PTEN*).

#### Fixed tissue Immunohistochemistry and imaging

For immunohistochemistry, CNS from staged larvae were dissected in PBS, fixed for 20 min in 4% formaldehyde diluted in PBS, washed three times in PBS-T (PBS+0.3% Triton X-100) and incubated two nights at 4°C with primary antibodies diluted in blocking solution (PBS-T, 5% Bovine Serum Albine, 2% Normal Goat Serum). After washing three times in PBS-T, CNS were incubated overnight at 4°C or 3-3 h at room temperature with secondary antibodies (dilution 1:200) diluted in blocking solution. Brains were washed three times in PBS-T and mounted in Mowiol mounting medium on a borosilicate glass side (number 1.5; VWR International). For the Nrx-IV antibody, CNS were fixed for 3 minutes in Bouin’s fixative solution (Sigma Aldrich, HT10132), and the rest of the protocol was identical. Primary antibodies used were: guinea pig anti-Dpn (1:5000,[25]), chicken anti-GFP (1:2000, Abcam ab13970), rat anti-ELAV (1:100, 7E8A10-c, DSHB), mouse anti-ELAV (1:100, 9F8A9-c, DSHB), rat anti-dE-cadherin (1:50, DCAD2, DSHB), mouse anti-Armadillo (1:50, N2 7A1, DSHB), rabbit anti-Repo (1:10000, kind gift from B. Altenheim), mouse anti-Repo 1:100 (DSHB, 8D12-c), mouse anti-Prospero (1:100, MR1A, DSHB), rabbit anti-Asense (1:3000, kind gift from the Yan lab). rabbit anti-Phospho-histone H3 (1:100, Millipore 06-570), rabbit anti-Nrx-IV (1:1000, [46]) mouse anti-wrapper (1:20, DSHB 10D3, supernatant), mouse anti-Nrg^180^ (1:50, DSHB BP104, supernatant). Fluorescently-conjugated secondary antibodies Alexa Fluor 405, Alexa Fluor 488, Alexa Fluor 546 and Alexa Fluor 633 (ThermoFisher Scientific) were used at a 1:200 dilution. DAPI (4′,6-diamidino-2-phenylindole, ThermoFisher Scientific 62247) was used to counterstain the nuclei.

#### Image acquisition and processing

Confocal images were acquired using a laser scanning confocal microscope (Zeiss LSM 880, Zen software (2012 S4)) with a Plan-Apochromat 40x/1.3 Oil objective. All brains were imaged as z-stacks with each section corresponding to 0.3-0.5 μm. The spectral mode was used for acquiring pictures of Raeppli clones. Images were subsequently analysed and processed using Fiji (Schindelin, J. 2012), Volocity (6.3 Quorum technologies), and the Open-Source software Icy v2.1.4.0 (Institut Pasteur and France Bioimaging, license GPLv3). .Denoising was used for some images using the Remove noise function (Fine filter) in Volocity. Images were assembled using Adobe Illustrator 25.4.6.

#### Multicolour clonal analyses (Raeppli)

Raeppli clones were generated by subjected freshly hatched larvae (ALH0) to a 37°C heat shocked for 2 hours. The genetic crosses and culture conditions were the following:

• Fig. 1G:

*yw, hs-FLP; Nrv2∷GFP, wor-GAL4/CyO*

x *UAS-Raeppli-NLS 53D; UAS-pros RNAi, tubGAL80^ts^*

x *UAS-Raeppli-NLS 53D; UAS-brat RNAi, tubGAL80^ts^*

x *UAS-Raeppli-NLS 53D; tubGAL80^ts^*

For *brat* RNAI, larvae were subjected to a 2 h heatshock at 37°C just after larval haching (ALH0), then transferred to 29°C.

For *pros* RNAi, larvae were kept at 18°C for 48h after collection and then subjected to a 2 h heatshock at 37°C. The larvae were transferred to 29°C afterwards.

• Fig. 2B, 3E and Supp. Fig. 2B, 3F-G:

*tub-QS; Nrv2∷GFP, wor-GAL4/CyO*

x *yw, hs-FLP; UAS-Raeppli-NLS 53D*

x *yw, hs-FLP; UAS-Raeppli-NLS 53D, QUAS-PTEN*

x *yw, hs-FLP; UAS-Raeppli-NLS 53D, QUAS-PTEN; UAS-shg RNAi VDRC 27082*

72h at 29°C on plates with 20 mg/ml quinic acid (T1) followed by 28h at 29°C on plates without quinic acid (T2)

• Fig. 3D:

*Nrv2∷GFP; wor-GAL4/CyO;*

x *yw, hs-FLP; UAS-Raeppli-nls 53D; UAS-shg RNAi VDRC 27082*

Larvae were kept 68h at 29°C from ALH0.

• Fig. 4D:

*Nrv2∷GFP; wor-GAL4/CyO; UAS-Raeppli-nls 89A*

x yw, *hs-FLP; UAS-Nrx-IV RNAi*

Larvae were kept 72h at 29°C from ALH0.

• Fig. 4G :

*yw, hs-FLP; Nrv2∷GFP; wor-GAL4/CyO; tub-Gal80^ts^*

x *UAS-nrg RNAi; UAS-Raeppli-NLS 89A*

Just hatched larvae (ALH0) were subjected to a 2 h heatshock at 37°C and then kept at 18°C for 24 h after collection. The larvae were transferred to 29°C afterwards, and dissected 54 h later.

• Fig. 6B:

*yw, hs-FLP; Nrv2∷GFP; wor-GAL4/CyO; tub-Gal80^ts^*

x *UAS-Raeppli-CAAX 42D*

x *UAS-Raeppli-CAAX 42D; UAS-Nrx-IV RNAi*

Just hatched larvae (ALH0) were subjected to a 2 h heatshock at 37°C, transferred to 29°C afterwards, and dissected 72 h later.

• • Fig. 6C:

yw, *hs-FLP; Nrv2∷GFP, wor-GAL4/CyO; tub-GAL80^ts^*

x *UAS-Raeppli-CAAX 99E*

x *UAS-nrg RNAi, UAS-Raeppli-CAAX 99E*

Just hatched larvae (ALH0) were subjected to a 2 h heatshock at 37°C and then kept at 18°C for 24 h after collection. The larvae were transferred to 29°C afterwards, and dissected 54 h later.

#### *shg^null^* MARCM clones

*shg^64R^, FRT42B; cyp4g15-myr∷dTomato* flies were crossed to the TUG13 MARCM line (*y,w, hs-FLP; FRTG13, tubP-GAL80[LL2]/ (CyO, act-GFP[JMR1]); tubP-GAL4[LL7], UAS-mCD8-GFP[LL6]/TM6B*). The resulting progeny was let to develop at 25°C, then subjected to 37°C heatshock either at 14-18 h after egg laying for 2 h, or at ALH48 for 30 min, and finally dissected at ALH72 (Supp. Fig. 3C).

#### Clonal analyses using CoinFLP

The Coin-FLP method [36] was used to induce rare clones of PTEN expressing CG cells, by crossing *cyp4g15-FLP; CoinFLP GAL4∷LexA; UAS-mCD8∷RFP* females to to *UAS-PTEN* males or *w1118* males for control, and maintained at 25°C. Larvae were staged to ALH48-ALH72 at 25°C.

#### Quantification of NSC lineage encasing (Raeppli-NLS)

For each VNC the total number of chambers containing more than one NSC, and the corresponding total number of NSC lineages non-individually encased were determined by counting manually the number of different Raeppli clones (colours) contained within one continuous CG membrane, choosing the z plane where differentiating lineages were well visible (at mid-disance between the CNS surface and the neuropile).

#### Quantification of NSC mitotic index

CNSs of the chosen genotypes were stained with Dpn and phospho-histone H3 antibodies to detect NSC fate and mitotis, respectively. Quantification was performed on NSCs from the thoracic part of the VNC. Mitotic phases (Prophase, Prometaphase/metaphase, Anaphase, Telophase) were manually determined by the localization and pattern of PH3^+^ DNA and Dpn staining. Normalized mitotic index corresponds to the ratio between mitotic NSCs over all NSCs, then divided by the mean of this ratio for the control sample.

#### Quantification of axonal angle

Z stacks of Raeppli CAAX clones induced in NSC lineages were visualized in Volocity (6.3 Quorum technologies), in a lateral view along the antero-posterior axis (xz axis). The brightest colour (mTFP1) was chosen and lines were drawn parallel to and following the main axonal projection for each clone. Pictures of the line were recorded through snapshots, and imported into Icy v2.1.4.0 where the Angle Helper plugin was used to measure the angle formed with the intersection with the AP axis (Fig. 6D-F).

#### Behavioural analysis

Locomotor activity of individual flies was measured with the *Drosophila* ethoscope activity Open Source system [60] at 23°C. Young adult males (7-10 days) were individually placed in 6.5 cm transparent tubes containing 1.5 cm nutrient medium (agarose 2% (p/v), sucrose 5% (p/v)) closed with wax at one end and cotton at the other. 20 tubes were positioned in each ethoscope and flies were first entrained to 12hr:12hr light-dark (LD) cycles for 3 days and their activity was recorded for 3 more days. The activity data analysis was done with the R software, using Rethomics packages [74]. Flies were considered asleep when found motionless for at least 5 min. The percentage of time sleeping across LD cycles was measured as the fraction of time sleep within 30 min intervals, whereas global time sleeping per fly corresponds to a total sleep over total time ratio. Velocity was measured using a previously described tracking algorithm [74] and is expressed in relative units. Fly moving across LD cycles was calculated as the fraction of time moving within 30 min time intervals. Overall, 35 control (non induced) flies and 55 (induced) flies, coming from 8 independent experiments, were analysed for each genotype. Genotypes positions in 20 tubes arenas were changed from one experiment to another to avoid positional bias.

#### Statistics and reproducibility

Statistical tests used for each experiment are stated in the figure legends. Statistical tests were performed using GraphPad Prism 7.0a. For all box and whisker plots, whiskers mark the minimum and maximum, the box includes the 25th–75th percentile, and the line in the box is the median. Individual values are superimposed.

## Acknowledgments

We thank Y. Bellaïche, A. Brand, Y-N Jan, M. Landgraf, R. Le Borgne, C. Klämbt and C. Maurange for reagents and strains. L. Arbogast generated the *cyp4g15-FRT-STOP-FRT-GAL4* and QUAS-PTEN constructs. Stocks obtained from the Bloomington Drosophila Stock Center (NIH P40OD018537) and from the Vienna Drosophila Resource Center were used in this study. Antibodies were obtained for the Developmental Studie Hybridoma Bank. This work has been funded by a starting package from Institut Pasteur/ LabEx Revive, a JCJC grant from Agence Nationale de la Recherche (NeuraSteNic, ANR-17-CE13-0010-01) to P.S. and a Projet Fondation ARC from the Association pour la Recherche contre le Cancer to P.S. A. B-L. was supported by a post-doctoral fellowship from the LabEx Revive.

## Author Contributions

A.B-L, P.S., V.R and I.G. conceived the experimental design. A.B-L. performed experiments of Figs. 1; 2; 3A-C, F-G and Supp. Figs. 1B-C; 2; 3A-B, H. P.S. performed experiments of Figs. 3D, H; 4C-H; 5; 6 and Supp. Fig. 3C,D,G; 44; 5; 6. V.R. performed experiments of Figs. 7A-C and Supp. Fig. 7. D.B. performed experiments of Fig 4A-B. A.B-L., V.R. and P.S. analyzed the data. A.B-L., V.R. and P.S. wrote the manuscript.

## Declaration of Interests

The authors declare no competing interest.

## Data availability statement

The datasets generated during and/or analysed during the current study are available from the corresponding author on reasonable request.

## Notes

### Competing Interest Statement

The authors have declared no competing interest.

## References

1. Ferraro, F., Lo Celso, C., and Scadden, D. (2010). Adult stem cells and their niches. Adv. Exp. Med. Biol. 695, 155–168. Available at: http://www.ncbi.nlm.nih.gov/pubmed/21222205 [Accessed May 12, 2020].

2. Lander, A.D., Kimble, J., Clevers, H., Fuchs, E., Montarras, D., Buckingham, M., Calof, A.L., Trumpp, A., and Oskarsson, T. (2012). What does the concept of the stem cell niche really mean today? BMC Biol. 10, 19. Available at: http://www.pubmedcentral.nih.gov/articlerender.fcgi?artid=3298504&tool=pmcentrez&rendertype=abstract [Accessed April 3, 2016].

3. Chacón-Martínez, C.A., Koester, J., and Wickström, S.A. (2018). Signaling in the stem cell niche: regulating cell fate, function and plasticity. Development 145, dev165399.

4. Decimo, I., Bifari, F., Krampera, M., and Fumagalli, G. (2012). Neural stem cell niches in health and diseases. Curr. Pharm. Des. 18, 1755–83. Available at: http://www.ncbi.nlm.nih.gov/pubmed/22394166 [Accessed October 14, 2019].

5. Obernier, K., and Alvarez-Buylla, A. (2019). Neural stem cells: Origin, heterogeneity and regulation in the adult mammalian brain. Dev. 146.

6. Randolph S. Ashton, Anthony Conway, Chinmay Pangarkar, J.B., and Kwang-Il Lim, Priya Shah, Mina Bissell, and D.V.S. (2012). Astrocytes regulate adult hippocampal neurogenesis through ephrin-B signaling. Nat. Neurosci. 9, 47–55.

7. Paul, A., Chaker, Z., and Doetsch, F. (2017). Hypothalamic regulation of regionally distinct adult neural stem cells and neurogenesis. Science (80-.). 356, 1383–1386. Available at: http://www.sciencemag.org/lookup/doi/10.1126/science.aal3839.

8. Chell, J.M., and Brand, A.H. (2010). Nutrition-responsive glia control exit of neural stem cells from quiescence. Cell 143, 1161–73. Available at: http://www.pubmedcentral.nih.gov/articlerender.fcgi?artid=3087489&tool=pmcentrez&rendertype=abstract [Accessed February 27, 2013].

9. Sousa-Nunes, R., Yee, L.L., and Gould, A.P. (2011). Fat cells reactivate quiescent neuroblasts via TOR and glial insulin relays in Drosophila. Nature 471, 508–12. Available at: http://www.pubmedcentral.nih.gov/articlerender.fcgi?artid=3146047&tool=pmcentrez&rendertype=abstract [Accessed February 27, 2013].

10. Homem, C.C.F., Repic, M., and Knoblich, J.A. (2015). Proliferation control in neural stem and progenitor cells. Nat. Rev. Neurosci. 16, 647–59. Available at: http://www.ncbi.nlm.nih.gov/pubmed/26420377 [Accessed April 12, 2016].

11. Bello, B.C., Izergina, N., Caussinus, E., and Reichert, H. (2008). Amplification of neural stem cell proliferation by intermediate progenitor cells in Drosophila brain development. Neural Dev. 3, 5. Available at: http://www.pubmedcentral.nih.gov/articlerender.fcgi?artid=2265709&tool=pmcentrez&rendertype=abstract [Accessed March 1, 2013].

12. Boone, J.Q., and Doe, C.Q. (2008). Identification of Drosophila type II neuroblast lineages containing transit amplifying ganglion mother cells. Dev. Neurobiol. 68, 1185–95. Available at: http://www.pubmedcentral.nih.gov/articlerender.fcgi?artid=2804867&tool=pmcentrez&rendertype=abstract [Accessed March 4, 2013].

13. Bowman, S.K., Rolland, V., Betschinger, J., Kinsey, K. a, Emery, G., and Knoblich, J. a (2008). The tumor suppressors Brat and Numb regulate transit-amplifying neuroblast lineages in Drosophila. Dev. Cell 14, 535–46. Available at: http://www.pubmedcentral.nih.gov/articlerender.fcgi?artid=2988195&tool=pmcentrez&rendertype=abstract [Accessed March 1, 2013].

14. Spéder, P., and Brand, A.H. (2014). Gap Junction Proteins in the Blood-Brain Barrier Control Nutrient-Dependent Reactivation of Drosophila Neural Stem Cells. Dev. Cell, 309–321. Available at: http://www.ncbi.nlm.nih.gov/pubmed/25065772 [Accessed August 1, 2014].

15. Kanai, M.I., Kim, M.-J., Akiyama, T., Takemura, M., Wharton, K., O’Connor, M.B., and Nakato, H. (2018). Regulation of neuroblast proliferation by surface glia in the Drosophila larval brain. Sci. Rep. 8, 3730. Available at: http://www.nature.com/articles/s41598-018-22028-y [Accessed May 24, 2019].

16. Morante, J., Vallejo, D.M., Desplan, C., and Dominguez, M. (2013). Conserved miR-8/miR-200 defines a glial niche that controls neuroepithelial expansion and neuroblast transition. Dev. Cell 27, 174–187. Available at: https://pubmed.ncbi.nlm.nih.gov/24139822/ [Accessed August 5, 2022].

17. Plazaola-Sasieta, H., Zhu, Q., Gaitán-Peñas, H., Rios, M., Estévez, R., and Morey, M. (2019). Drosophila ClC-a is required in glia of the stem cell niche for proper neurogenesis and wiring of neural circuits. Glia 67, 2374–2398. Available at: https://onlinelibrary.wiley.com/doi/full/10.1002/glia.23691 [Accessed April 7, 2021].

18. Dong, Q., Zavortink, M., Froldi, F., Golenkina, S., Lam, T., and Cheng, L.Y. (2021). Glial Hedgehog signalling and lipid metabolism regulate neural stem cell proliferation in Drosophila. EMBO Rep. Available at: https://pubmed.ncbi.nlm.nih.gov/33751817/ [Accessed April 18, 2021].

19. Cheng, L.Y., Bailey, A.P., Leevers, S.J., Ragan, T.J., Driscoll, P.C., and Gould, A.P. (2011). Anaplastic lymphoma kinase spares organ growth during nutrient restriction in Drosophila. Cell 146, 435–47. Available at: http://www.ncbi.nlm.nih.gov/pubmed/21816278 [Accessed February 27, 2013].

20. Bailey, A.P., Koster, G., Guillermier, C., Hirst, E.M.A., MacRae, J.I., Lechene, C.P., Postle, A.D., and Gould, A.P. (2015). Antioxidant Role for Lipid Droplets in a Stem Cell Niche of Drosophila. Cell 163, 340–353. Available at: http://www.pubmedcentral.nih.gov/articlerender.fcgi?artid=4601084&tool=pmcentrez&rendertype=abstract [Accessed October 9, 2015].

21. Spéder, P., and Brand, A.H. (2018). Systemic and local cues drive neural stem cell niche remodelling during neurogenesis in Drosophila. Elife 7, e30413. Available at: https://elifesciences.org/articles/30413 [Accessed July 8, 2018].

22. Dumstrei, K., Wang, F., and Hartenstein, V. (2003). Role of DE-cadherin in neuroblast proliferation, neural morphogenesis, and axon tract formation in Drosophila larval brain development. J. Neurosci. 23, 3325–35. Available at: http://www.ncbi.nlm.nih.gov/pubmed/12716940.

23. Pereanu, W., Shy, D., and Hartenstein, V. (2005). Morphogenesis and proliferation of the larval brain glia in Drosophila. Dev. Biol. 283, 191–203. Available at: http://www.ncbi.nlm.nih.gov/pubmed/15907832 [Accessed March 1, 2013].

24. Yuan, X., Sipe, C.W., Suzawa, M., Bland, M.L., and Siegrist, S.E. (2020). Dilp-2-mediated PI3-kinase activation coordinates reactivation of quiescent neuroblasts with growth of their glial stem cell niche. PLoS Biol. 18, 1–24. Available at: http://dx.doi.org/10.1371/journal.pbio.3000721.

25. Rujano, M.A., Briand, D., Ðelić, B., Marc, J., and Spéder, P. (2022). An interplay between cellular growth and atypical fusion defines morphogenesis of a modular glial niche in Drosophila. Nat. Commun. 13. Available at: https://pubmed.ncbi.nlm.nih.gov/36008397/ [Accessed September 26, 2022].

26. Spindler, S.R., and Hartenstein, V. (2010). The Drosophila neural lineages: a model system to study brain development and circuitry. Dev. Genes Evol. 220, 1–10. Available at: http://www.ncbi.nlm.nih.gov/pubmed/20306203 [Accessed June 21, 2018].

27. Larsen, C., Shy, D., Spindler, S.R., Fung, S., Pereanu, W., Younossi-Hartenstein, A., and Hartenstein, V. (2009). Patterns of growth, axonal extension and axonal arborization of neuronal lineages in the developing Drosophila brain. Dev. Biol. 335, 289. Available at: pmc/articles/PMC2785225/ [Accessed September 29, 2022].

28. Spindler, S.R., Ortiz, I., Fung, S., Takashima, S., and Hartenstein, V. (2009). Drosophila cortex and neuropile glia influence secondary axon tract growth, pathfinding, and fasciculation in the developing larval brain. Dev. Biol. 334, 355–368.

29. Freeman, M.R. (2015). Drosophila Central Nervous System Glia. Cold Spring Harb. Perspect. Biol. 7. Available at: http://www.ncbi.nlm.nih.gov/pubmed/25722465 [Accessed June 14, 2018].

30. Kang, K.H., and Reichert, H. (2015). Control of neural stem cell self-renewal and differentiation in Drosophila. Cell Tissue Res. 359, 33–45. Available at: http://www.ncbi.nlm.nih.gov/pubmed/24902665 [Accessed October 30, 2015].

31. Choksi, S.P., Southall, T.D., Bossing, T., Edoff, K., de Wit, E., Fischer, B.E., van Steensel, B., Micklem, G., and Brand, A.H. (2006). Prospero acts as a binary switch between self-renewal and differentiation in Drosophila neural stem cells. Dev Cell 11, 775–789. Available at: http://www.ncbi.nlm.nih.gov/entrez/query.fcgi?cmd=Retrieve&db=PubMed&dopt=Citation&list_uids=17141154.

32. Bello, B., Reichert, H., and Hirth, F. (2006). The brain tumor gene negatively regulates neural progenitor cell proliferation in the larval central brain of Drosophila. Development 133, 2639–2648.

33. Betschinger, J., Mechtler, K., and Knoblich, J. a (2006). Asymmetric segregation of the tumor suppressor brat regulates self-renewal in Drosophila neural stem cells. Cell 124, 1241–1253. Available at: http://www.ncbi.nlm.nih.gov/pubmed/16564014 [Accessed February 27, 2013].

34. Kanca, O., Caussinus, E., Denes, A.S., Percival-Smith, A., and Affolter, M. (2014). Raeppli: a whole-tissue labeling tool for live imaging of Drosophila development. Development 141, 472–480. Available at: http://www.ncbi.nlm.nih.gov/pubmed/24335257.

35. Foty, R.A., and Steinberg, M.S. (2013). Differential adhesion in model systems. Wiley Interdiscip. Rev. Dev. Biol. 2, 631–645. Available at: https://onlinelibrary.wiley.com/doi/full/10.1002/wdev.104 [Accessed September 5, 2022].

36. Bosch, J.A., Tran, N.H., and Hariharan, I.K. (2015). CoinFLP: a system for efficient mosaic screening and for visualizing clonal boundaries in Drosophila. Development 142, 597–606. Available at: http://dev.biologists.org/cgi/doi/10.1242/dev.114603.

37. White, K., Grether, M.E., Abrams, J.M., Young, L., Farrell, K., and Steller, H. (1994). Genetic control of programmed cell death in Drosophila. Science (80-.). 264, 677–683. Available at: http://www.ncbi.nlm.nih.gov/entrez/query.fcgi?cmd=Retrieve&db=PubMed&dopt=Citation&list_uids=8171319.

38. Coutinho-Budd, J.C., Sheehan, A.E., and Freeman, M.R. (2017). The secreted neurotrophin spätzle 3 promotes glial morphogenesis and supports neuronal survival and function. Genes Dev. 31, 2023–2038.

39. Almeida, M.S., and Bray, S.J. (2005). Regulation of post-embryonic neuroblasts by Drosophila Grainyhead. Mech. Dev. 122, 1282–1293.

40. Nériec, N., and Desplan, C. (2016). From the Eye to the Brain: Development of the Drosophila Visual System. Curr. Top. Dev. Biol. 116, 247–71. Available at: http://www.ncbi.nlm.nih.gov/pubmed/26970623 [Accessed April 4, 2016].

41. Doyle, S.E., Pahl, M.C., Siller, K.H., Ardiff, L., and Siegrist, S.E. (2017). Neuroblast niche position is controlled by phosphoinositide 3-kinase-dependent DE-cadherin adhesion. Dev. 144, 820–829.

42. Hortsch, M., and Margolis, B. (2003). Septate and paranodal junctions: kissing cousins. Trends Cell Biol. 13, 557–561.

43. Zihni, C., Mills, C., Matter, K., and Balda, M.S. (2016). Tight junctions: from simple barriers to multifunctional molecular gates. Nat. Rev. Mol. Cell Biol. 2016 179 17, 564–580. Available at: https://www.nature.com/articles/nrm.2016.80 [Accessed September 30, 2022].

44. Baumgartner, S., Littleton, J.T., Broadie, K., Bhat, M. a, Harbecke, R., Lengyel, J. a, Chiquet-Ehrismann, R., Prokop, a, and Bellen, H.J. (1996). A Drosophila neurexin is required for septate junction and blood-nerve barrier formation and function. Cell 87, 1059–68. Available at: http://www.ncbi.nlm.nih.gov/pubmed/8978610.

45. Noordermeer, J.N., Kopczynski, C.C., Fetter, R.D., Bland, K.S., Chen, W.Y., and Goodman, C.S. (1998). Wrapper, a novel member of the Ig superfamily, is expressed by midline glia and is required for them to ensheath commissural axons in Drosophila. Neuron 21, 991–1001. Available at: https://pubmed.ncbi.nlm.nih.gov/9856456/ [Accessed August 29, 2022].

46. Stork, T., Thomas, S., Rodrigues, F., Silies, M., Naffin, E., Wenderdel, S., and Klämbt, C. (2009). Drosophila Neurexin IV stabilizes neuron-glia interactions at the CNS midline by binding to Wrapper. Development 136, 1251–61. Available at: http://www.ncbi.nlm.nih.gov/pubmed/19261699 [Accessed March 4, 2013].

47. Wheeler, S.R., Banerjee, S., Blauth, K., Rogers, S.L., Bhat, M.A., and Crews, S.T. (2009). Neurexin IV and Wrapper interactions mediate Drosophila midline glial migration and axonal ensheathment. Development 136, 1147–1157. Available at: https://journals.biologists.com/dev/article/136/7/1147/65448/Neurexin-IV-and-Wrapper-interactions-mediate [Accessed September 6, 2022].

48. Yamamoto, M., Ueda, R., Takahashi, K., Saigo, K., and Uemura, T. (2006). Control of Axonal Sprouting and Dendrite Branching by the Nrg-Ank Complex at the Neuron-Glia Interface. Curr. Biol. 16, 1678–1683.

49. Martin, V., Mrkusich, E., Steinel, M.C., Rice, J., Merritt, D.J., and Whitington, P.M. (2008). The L1-type cell adhesion molecule Neuroglian is necessary for maintenance of sensory axon advance in the Drosophila embryo. Neural Dev. 3, 1–11. Available at: https://neuraldevelopment.biomedcentral.com/articles/10.1186/1749-8104-3-10 [Accessed September 6, 2022].

50. Yang, W.K., Chueh, Y.R., Cheng, Y.J., Siegenthaler, D., Pielage, J., and Chien, C.T. (2019). Epidermis-Derived L1CAM Homolog Neuroglian Mediates Dendrite Enclosure and Blocks Heteroneuronal Dendrite Bundling. Curr. Biol. 29, 1445–1459.e3.

51. Goossens, T., Kang, Y.Y., Wuytens, G., Zimmermann, P., Callaerts-Végh, Z., Pollarolo, G., Islam, R., Hortsch, M., and Callaerts, P. (2011). The Drosophila L1CAM homolog Neuroglian signals through distinct pathways to control different aspects of mushroom body axon development. Development 138, 1595–1605. Available at: https://pubmed.ncbi.nlm.nih.gov/21389050/ [Accessed September 9, 2022].

52. Clements, J., Buhler, K., Winant, M., Vulsteke, V., and Callaerts, P. (2021). Glial and Neuronal Neuroglian, Semaphorin-1a and Plexin A Regulate Morphological and Functional Differentiation of Drosophila Insulin-Producing Cells. Front. Endocrinol. (Lausanne). 12. Available at: https://pubmed.ncbi.nlm.nih.gov/34276554/ [Accessed September 30, 2022].

53. Shepherd, D., Harris, R., Williams, D.W., and Truman, J.W. (2016). Postembryonic lineages of the Drosophila ventral nervous system: Neuroglian expression reveals the adult hemilineage associated fiber tracts in the adult thoracic neuromeres. J. Comp. Neurol. 524, 2677–2695. Available at: https://onlinelibrary.wiley.com/doi/full/10.1002/cne.23988 [Accessed September 9, 2022].

54. Nakano, R., Iwamura, M., Obikawa, A., Togane, Y., Hara, Y., Fukuhara, T., Tomaru, M., Takano-Shimizu, T., and Tsujimura, H. (2019). Cortex glia clear dead young neurons via Drpr/dCed-6/Shark and Crk/Mbc/dCed-12 signaling pathways in the developing Drosophila optic lobe. Dev. Biol. Available at: https://www.sciencedirect.com/science/article/pii/S0012160618305645?via%3Dihub [Accessed May 13, 2019].

55. Lee, P.T., Zirin, J., Kanca, O., Lin, W.W., Schulze, K.L., Li-Kroeger, D., Tao, R., Devereaux, C., Hu, Y., Chung, V., et al. (2018). A gene-specific T2A-GAL4 library for drosophila. Elife 7.

56. Hortsch, M., Bieber, A.J., Patel, N.H., and Goodman, C.S. (1990). Differential splicing generates a nervous system-specific form of Drosophila neuroglian. Neuron 4, 697–709. Available at: https://pubmed.ncbi.nlm.nih.gov/1693086/ [Accessed September 13, 2022].

57. Hortsch, M., Bieber, A.J., Patel, N.H., and Goodman, C.S. (1990). Differential splicing generates a nervous system—Specific form of drosophila neuroglian. Neuron 4, 697–709.

58. Islam, R., Kristiansen, L. V., Romani, S., Garcia-Alonso, L., and Hortsch, M. (2004). Activation of EGF receptor kinase by L1-mediated homophilic cell interactions. Mol. Biol. Cell 15, 2003–2012. Available at: https://pubmed.ncbi.nlm.nih.gov/14718570/ [Accessed September 20, 2022].

59. Goossens, T., Kang, Y.Y., Wuytens, G., Zimmermann, P., Callaerts-Végh, Z., Pollarolo, G., Islam, R., Hortsch, M., and Callaerts, P. (2011). The Drosophila L1CAM homolog Neuroglian signals through distinct pathways to control different aspects of mushroom body axon development. Development 138, 1595–1605. Available at: https://pubmed.ncbi.nlm.nih.gov/21389050/ [Accessed September 6, 2022].

60. Geissmann, Q., Garcia Rodriguez, L., Beckwith, E.J., French, A.S., Jamasb, A.R., and Gilestro, G.F. (2017). Ethoscopes: An open platform for high-throughput ethomics. PLOS Biol. 15, e2003026. Available at: https://journals.plos.org/plosbiology/article?id=10.1371/journal.pbio.2003026 [Accessed September 15, 2022].

61. Beckwith, E.J., and French, A.S. (2019). Sleep in Drosophila and Its Context. Front. Physiol. 10, 1167. Available at: pmc/articles/PMC6749028/ [Accessed February 22, 2022].

62. Kashima, R., Redmond, P.L., Ghatpande, P., Roy, S., Kornberg, T.B., Hanke, T., Knapp, S., Lagna, G., and Hata, A. (2017). Hyperactive locomotion in a Drosophila model is a functional readout for the synaptic abnormalities underlying fragile X syndrome. Sci. Signal. 10. Available at: https://www.science.org/doi/10.1126/scisignal.aai8133 [Accessed September 30, 2022].

63. Klein, M., Singgih, E.L., van Rens, A., Demontis, D., Børglum, A.D., Mota, N.R., Castells-Nobau, A., Kiemeney, L.A., Brunner, H.G., Arias-Vasquez, A., et al. (2020). Contribution of Intellectual Disability-Related Genes to ADHD Risk and to Locomotor Activity in Drosophila. Am. J. Psychiatry 177, 526–536. Available at: https://pubmed.ncbi.nlm.nih.gov/32046534/ [Accessed September 30, 2022].

64. Takai, A., Chiyonobu, T., Ueoka, I., Tanaka, R., Tozawa, T., Yoshida, H., Morimoto, M., Hosoi, H., and Yamaguchi, M. (2020). A novel Drosophila model for neurodevelopmental disorders associated with Shwachman–Diamond syndrome. Neurosci. Lett. 739, 135449.

65. Yang, Z., Yu, Y., Zhang, V., Tian, Y., Qi, W., and Wang, L. (2015). Octopamine mediates starvation-induced hyperactivity in adult Drosophila. Proc. Natl. Acad. Sci. U. S. A. 112, 5219–5224. Available at: https://www.pnas.org/doi/abs/10.1073/pnas.1417838112 [Accessed September 30, 2022].

66. Yu, Y., Huang, R., Ye, J., Zhang, V., Wu, C., Cheng, G., Jia, J., and Wang, L. (2016). Regulation of starvation-induced hyperactivity by insulin and glucagon signaling in adult Drosophila. Elife 5.

67. Edenfeld, G., Volohonsky, G., Krukkert, K., Naffin, E., Lammel, U., Grimm, A., Engelen, D., Reuveny, A., Volk, T., Klambt, C., et al. (2006). The splicing factor crooked neck associates with the RNA-binding protein HOW to control glial cell maturation in Drosophila. Neuron 52, 969–980. Available at: http://www.ncbi.nlm.nih.gov/pubmed/17178401 [Accessed March 10, 2013].

68. Zhu, S., Barshow, S., Wildonger, J., Jan, L.Y., and Jan, Y.-N. (2011). Ets transcription factor Pointed promotes the generation of intermediate neural progenitors in Drosophila larval brains. Proc. Natl. Acad. Sci. U. S. A. 108, 20615–20. Available at: http://www.ncbi.nlm.nih.gov/pubmed/22143802 [Accessed October 4, 2022].

69. Sotillos, S., Díaz-Meco, M.T., Caminero, E., Moscat, J., and Campuzano, S. (2004). DaPKC-dependent phosphorylation of Crumbs is required for epithelial cell polarity in Drosophila. J. Cell Biol. 166, 549–557. Available at: http://www.jcb.org/cgi/doi/10.1083/jcb.200311031 [Accessed September 26, 2022].

70. Pfeiffer, B.D., Jenett, A., Hammonds, A.S., Ngo, T.-T.B., Misra, S., Murphy, C., Scully, A., Carlson, J.W., Wan, K.H., Laverty, T.R., et al. (2008). Tools for neuroanatomy and neurogenetics in Drosophila. Proc. Natl. Acad. Sci. U. S. A. 105, 9715–20. Available at: http://www.pubmedcentral.nih.gov/articlerender.fcgi?artid=2447866&tool=pmcentrez&rendertype=abstract.

71. Petersen, L.K., and Stowers, R.S. (2011). A Gateway MultiSite recombination cloning toolkit. PLoS One 6, e24531. Available at: http://www.pubmedcentral.nih.gov/articlerender.fcgi?artid=3170369&tool=pmcentrez&rendertype=abstract [Accessed March 9, 2016].

72. Huang, H., Potter, C.J., Tao, W., Li, D.M., Brogiolo, W., Hafen, E., Sun, H., and Xu, T. (1999). PTEN affects cell size, cell proliferation and apoptosis during Drosophila eye development. Development 126, 5365–5372. Available at: https://pubmed.ncbi.nlm.nih.gov/10556061/ [Accessed February 21, 2022].

73. Benmimoun, B., Papastefanaki, F., Périchon, B., Segklia, K., Roby, N., Miriagou, V., Schmitt, C., Dramsi, S., Matsas, R., and Spéder, P. (2020). An original infection model identifies host lipoprotein import as a route for blood-brain barrier crossing. Nat. Commun. 11, 1–18. Available at: https://doi.org/10.1038/s41467-020-19826-2 [Accessed March 9, 2021].

74. Geissmann, Q., Rodriguez, L.G., Beckwith, E.J., and Gilestro, G.F. (2019). Rethomics: An R framework to analyse high-throughput behavioural data. PLoS One 14. Available at: https://pubmed.ncbi.nlm.nih.gov/30650089/ [Accessed October 2, 2022].

