## Supplemental Figures for "A corset of adhesions during development establishes individual neural stem cell niches and controls adult behaviour"

**Supplemental Figure legends Banach-Latapy et al.**

**Supplemental Figure 1. Cortex glia individually encase normal and tumour NSC lineages.**

A) Confocal picture of one NSC lineage labelled with a membrane marker (mTFP1-CAAX, grey). CG membrane was visualized with *Nrv2::GFP* (green).

A') 3D reconstruction of the membrane signal of the NSC lineage shown in A).

B) Two different Type I tumours (*wor-GAL4 > aPKC*$^{CAAX}$ and *wor-GAL4 > Dpn*) were induced from ALH0 and the organisation of CG membrane was monitored by *Nrv2::GFP* (green). NSCs are labelled with Dpn (grey), and neuron with ElaV (magenta).

C) Representative confocal images of the progressive formation of individual encasing around Type II lineages, at ALH0, ALH16, ALH24, ALH30, ALH48, ALH72 and ALH96 at 25°C. CG membrane is visualized by *Nrv2::GFP* (green), NSCs are labelled with Dpn (grey), Type II NSCs are recognized by the absence of Ase (magenta) and GMCs are labelled with Pros (blue).

**Supplemental Figure 2. CG use intrinsic and generic NSC lineage cues to encase individual NSC lineages.**

A) Representative confocal pictures of the extent of individual encasing of NSC lineages by CG at T1, following the regimen described in Fig. 2A. Top panel shows the whole thoracic VNC, and bottom panel a close-up of the yellow box. NSC lineages were marked with the multicolour lineage tracing Raeppli-NLS (blue, white, orange and red), induced at ALH0 using *hs-Flp*. CG membrane was visualized with *Nrv2::GFP* (green). Dashed white lines outline zones where NSC lineages are still not individually encases. See Methods for timing, conditions and genetics of larval rearing.

B) Representative confocal picture of the CG network in control CG clone and in clone where apoptosis has been induced earlier (Reaper). The CoinFLP system was used to generate wild-type and Reaper clones in the CG. See Methods for timing, conditions and genetics of larval rearing. The membrane of the clone is marked by mCD8::RFP (magenta). CG membrane was visualized with *Nrv2::GFP* (green).

**Supplemental Figure 3. Intra-lineage adherens junctions are not absolutely required for individual encasing while providing robustness.**

A) Representative confocal pictures of the expression of Shg in Type I (*wor-GAL4 > pros RNAi*) and Type II (*PntP1-GAL4 > brat RNAi*) tumours. Organisation of CG membrane was monitored by *Nrv2::GFP* (green). Tumour NSCs are labelled with Dpn (grey), and Shg is detected with a specific antibody (magenta).

B) Representative confocal image of the expression of Shg in the optic lobe (ALH72 at 25°C). Shg is monitored through a *Shg::GFP* fusion (magenta), CG membrane is visualized by *Nrv2::GFP* (green), NSCs are labelled with Dpn (grey), and neurons are labelled with ElaV (blue).

C) Confocal images of mutant clones for *shg* in NSC lineages, induced during late embryogenesis (top panel) or at ALH48 at 25°C (bottom panel). The membrane of the clone is marked by mCD8::RFP (magenta). CG membrane is visualized with *Nrv2::GFP* (green), NSCs are labelled with Dpn (grey), Shg is detected with a specific antibody (blue, induction during embryogenesis) and neurons are labelled with ElaV (blue, induction at ALH48).

D) Representative confocal pictures of the extent of loss of signal for Shg (monitored through *Shg::GFP*, grey) in different genetic conditions defined by the driver line (CG, *cyp4g15-GAL4*; $L^{NSC}$, *wor-GAL4*) and RNAi constructs.

E) Representative confocal pictures of the thoracic VNC for *shg* knockdown by RNAi in the CG (*CG > shg RNAi*, driver line *Nrv2::GFP, tub-GAL80*$^{ts}$*; cyp4g15-GAL4;*). Larvae are dissected after 68 h at 29°C from ALH0. CG membrane is visualized by *Nrv2::GFP* (green) and NSCs are labelled with Dpn (grey).

F) Close-ups of confocal pictures assessing Shg levels in control, *shg* knockdown in NSC lineages (Fig. 3D) and *shg* knockdown in NSC lineages plus conditional block of CG growth (Fig. 3F). Shg levels are monitored with a specific antibody (magenta), NSC lineages are marked with the multicolour lineage tracing Raeppli-NLS (blue, white, orange and red) and CG membrane is visualized with *Nrv2::GFP* (green).

G) Quantification of the number of NSC lineages non-individually encased from Fig. 3D ($L^{NSC}$> *shg RNAi*). Control (n = 7 VNCs), and *shg RNAi* (n = 9 VNCs). Data statistics: Mann-Whitney test. Results are presented as box and whisker plots.

H) Representative confocal pictures of the extent of individual encasing of NSC lineages by CG at T1, following the regimen described in Fig. 3E. Top panel shows the whole thoracic VNC, and bottom panel a close-up of the yellow box. NSC lineages were marked with the multicolour lineage tracing Raeppli-NLS (blue, white, orange and red), induced at ALH0 using *hs-Flp*. CG membrane was visualized with *Nrv2::GFP* (green). See Methods for timing, conditions and genetics of larval rearing.

**Supplemental Figure 4. Neurexin-IV and Neuroglian are both required in NSC lineages for individual encasing.**

A) Representative confocal images of the expression of the septate junction components Dlg1, ATP$\alpha$ and Cora at ALH72 at 25°C. Dlg1 is monitored through a *Dlg1::GFP* fusion, ATP$\alpha$ through an *ATP$\alpha$::GFP* fusion and Cora is detected by a specific antibody (all in magenta). CG membrane is visualized by *Nrv2::GFP* (green), NSCs are labelled with Dpn (grey), and neurons are labelled with ElaV (blue).

B) Representative confocal pictures of the thoracic VNC for a condition in which *ATP$\alpha$* is knocked down by RNAi in NSC lineages ($L^{NSC}$ > *ATP$\alpha$ RNAi*, driver line *Nrv2::GFP, wor-GAL4; tub-GAL80$^{ts}$*). RNAi is induced after 3 days at 18°Cfrom ALH0, and larvae are dissected 48 h later at 29°C. CG membrane is visualized by *Nrv2::GFP* (green) and NSCs are labelled with Dpn (grey).

C) Representative confocal pictures of the thoracic VNC for control and for *nrx-IV* knockdown by RNAi in NSC lineages ($L^{NSC}$ > *nrx-IV RNAi*, driver line *Nrv2::GFP, wor-GAL4; tub-GAL80$^{ts}$*). Larvae are dissected after 68 h at 29°C from ALH0. Nrx-IV levels are monitored through *Nrx-IV::GFP* (magenta) and CG membrane is visualized by *Nrv2::GFP* (green).

D) Representative confocal pictures of the thoracic VNC for control and for *nrg* knockdown by RNAi in NSC lineages ($L^{NSC}$ > *nrg RNAi*, driver line *Nrv2::GFP, wor-GAL4; tub-GAL80$^{ts}$*). Larvae are kept 24 h at 18°C, then dissected after 54 h at 29°C. Nrg levels are monitored through *Nrg::GFP* (magenta) and NSCs are labelled with Dpn (grey).

E) Representative confocal pictures of thoracic VNCs for *nrx-IV* knockdown by RNAi in the neurons and in the CG (*ElaV-GAL4* and *cyp4g15-GAL4* drivers, respectively). Larvae are dissected after 68 h at 29°C from ALH0. CG membrane is visualized by *Nrv2::GFP* (green) and NSCs are labelled with Dpn (grey).

F) Representative confocal pictures of thoracic VNCs for *nrg* knockdown by RNAi in the neurons and in the CG (*ElaV-GAL4* and *cyp4g15-GAL4* drivers, respectively). For the neurons, larvae are kept 24 h at 18°C, then dissected after 54 h at 29°C.For the

CG, larvae are dissected after 68 h at 29°C from ALH0. CG membrane is visualized by *Nrv2::GFP* (green) and NSCs are labelled with Dpn (grey).

**Supplemental Figure 5. Individual encasing relies on balancing a CG to lineage interaction through Nrx-IV and Wrapper with an intra-lineage adhesion through Nrg.**

A) Representative confocal picture of the expression at ALH72 at 25°C of a CRIMIC line in which the GAL4 sequences are inserted in and under the control of *Wrapper* enhancers ($Wrapper^{CRIMIC}$). $Wrapper^{CRIMIC}$ drives fluorescent membrane (mCD8::GFP, green) and nuclear (Hist::RFP, yellow) reporters. NSCs are labelled with Dpn (grey) and glia with Repo (blue).

B) Representative confocal pictures of the expression of all Nrg isoforms (*Nrg::GFP*, green) compared to the $Nrg^{180}$ isoform (BP104, blue) at ALH72 at 25°C. Upper panel, top view. Lower panel, median cut though the NSC population. Note the absence of signal at the septate junctions (yellow arrowheads on the top view) for $Nrg^{180}$.

C) Representative confocal pictures and close-ups (C') of thoracic VNCs for control and for *nrg* knockdown by RNAi in NSC lineages ($L^{NSC}$ > *nrg RNAi*, driver line *Nrv2::GFP, wor-GAL4; tub-GAL80*$^{ts}$). Larvae are kept 24 h at 18°C, then dissected after 54 h at 29°C. Levels of the $Nrg^{180}$ isoform are monitored through staining with BP104 (blue). CG membrane is visualized by *Nrv2::GFP* (green) and NSCs are labelled with Dpn (grey).

D) Representative confocal picture of the respective localisations of the $Nrg^{167}$ and $Nrg^{180}$ isoform in a thoracic VNC, at ALH72 at 25°C. The $Nrg^{167}$ isoform is visualized by a protein trap in the *nrg* gene leading to the preferential expression of this isoform ($Nrg^{167}$*::GFP*, yellow). The $Nrg^{180}$ isoform is detected with a specific antibody (BP104, blue).

E) Representative confocal picture of the thoracic VNC for a condition in which $nrg^{GPI}$ (a transgene in which the transmembrane and cytoplasmic domains of Nrg are replaced by a GPI anchor signal) is overexpressed in the CG (*CG* > $nrg^{GPI}$, driver line *Nrv2::GFP, tub-GAL80*$^{ts}$*; cyp4g15-GAL4*). Larvae are dissected after 68 h at 29°C from ALH0. CG membrane is visualized by *Nrv2::GFP* (green) and NSCs are labelled with Dpn (grey).

F) Representative confocal picture of the thoracic VNC for a condition in which $nrg^{GPI}$ is overexpressed in NSC lineages ($L^{NSC}$ > $nrg^{GPI}$, driver line *Nrv2::GFP, wor-GAL4;*

*tub-GAL80^ts^*). Larvae are dissected after 68 h at 29°C from ALH0. CG membrane is visualized by *Nrv2::GFP* (green) and NSCs are labelled with Dpn (grey).

**Supplemental Figure 6. Loss of Nrx-IV and Nrg adhesions in NSC lineages during development does not alter NSC proliferation but induces axonal misprojection from newborn neurons.**

A) Mitotic index (left panel) and distribution of the mitotic phases (right panel) for control and *nrx-IV* knockdown in NSC lineages ($L^{NSC}$ > *nrx-IV RNAi,* driver line *Nrv2::GFP, wor-GAL4; tub-GAL80*$^{ts}$). Larvae are dissected after 68 h at 29°C from ALH0. Control (n = 10 VNCs) and *nrx-IV* RNAI (n = 9 VNCs). Data statistics: two-tailed unpaired t-test. Results are presented as box and whisker plots.

B) Mitotic index (left panel) and distribution of the mitotic phases (right panel) for control and *nrg* knockdown in NSC lineages ($L^{NSC}$ > *nrg RNAi,* driver line *Nrv2::GFP, wor-GAL4; tub-GAL80*$^{ts}$). Larvae are kept 24 h at 18°C, then dissected after 54 h at 29°C. Control (n = 8 VNCs) and *nrx-IV* RNAI (n = 10 VNCs). Data statistics: two-tailed unpaired t-test. Results are presented as box and whisker plots.

C) Mitotic index (left panel) and distribution of the mitotic phases (right panel) for control and *shg* knockdown in NSC lineages ($L^{NSC}$ > *shg RNAi,* driver line *Nrv2::GFP, wor-GAL4; tub-GAL80*$^{ts}$). Larvae are dissected after 68 h at 29°C from ALH0. Control (n = 10 VNCs) and *shg* RNAI (n = 7 VNCs). Data statistics: two-tailed unpaired t-test. Results are presented as box and whisker plots.

D) 3D reconstruction of a group of NSC lineages visualized with a membrane marker (mTFP1-CAAX) in antero-posterior view for *nrx-IV* knockdown in NSC lineages ($L^{NSC}$ > *nrx-IV RNAi*). Clonal labelling was obtained through the induction of Raeppli-CAAX in NSC lineages at ALH0. See Methods for timing, conditions and genetics of larval rearing. Colourings highlight aberrant projections.

**Supplemental Figure 7. Loss of Nrx-IV and Nrg adhesions in NSC lineages during development results in locomotor hyperactivity in the adults.**

A) Profiles of the averaged locomotor activity profiles over 24 h Light-Dark day for non-induced (flies always kept at 18°C before the recordings) and induced (flies shifted to 29°C from early larval stage to mid-pupal stage) conditions. Control, (*wor-GAL4, tub-Gal80*$^{ts}$ *x* $w^{1118}$), n = 35 non-induced adult males and n = 55 induced adult males. *shg* RNAi (*wor-GAL4, tub-Gal80*$^{ts}$ *x shg RNAi*$^{VDRC27082}$), n = 35 non-induced adult males and n = 55 induced adult males. *nrx-IV* RNAi (*wor-GAL4, tub-Gal80*$^{ts}$ *x nrx-IV RNAI*$^{BL32424}$), n = 35 non-induced adult males and n = 55 induced adult males. *nrg* RNAi (*wor-GAL4, tub-Gal80*$^{ts}$ *x nrg RNAI*$^{BL37496}$), n = 35 non-induced adult males and n = 55 induced adult males.

B) Percentage of global time sleeping during the Dark period (% asleep, ratio between total sleep time and total time) in non-induced (flies always kept at 18°C before the recordings) and induced (flies shifted to 29°C from early larval stage to mid-pupal stage) conditions. Control, (*wor-GAL4, tub-Gal80*$^{ts}$ *x w*$^{1118}$), n = 35 non-induced adult males and n = 55 induced adult males. *shg* RNAi (*wor-GAL4, tub-Gal80*$^{ts}$ *x shg RNAi*$^{VDRC27082}$), n = 35 non-induced adult males and n = 55 induced adult males. *nrx-IV* RNAi (*wor-GAL4, tub-Gal80*$^{ts}$ *x nrx-IV RNAI*$^{BL32424}$), n = 35 non-induced adult males and n = 55 induced adult males. *nrg* RNAi (*wor-GAL4, tub-Gal80*$^{ts}$ *x nrg RNAI*$^{BL37496}$), n = 35 non-induced adult males and n = 55 induced adult males. Data statistics: one-way ANOVA with a Kruskal–Wallis multiple comparison test. Results are presented as box and whisker plots.

C) Percentage of global time sleeping during the Light period (% asleep, ratio between total sleep time and total time) in non-induced (flies always kept at 18°C before the recordings) and induced (flies shifted to 29°C from early larval stage to mid-pupal stage) conditions. Control, (*wor-GAL4, tub-Gal80*$^{ts}$ *x w*$^{1118}$), n = 35 non-induced adult males and n = 55 induced adult males. *shg* RNAi (*wor-GAL4, tub-Gal80*$^{ts}$ *x shg RNAi*$^{VDRC27082}$), n = 35 non-induced adult males and n = 55 induced adult males. *nrx-IV* RNAi (*wor-GAL4, tub-Gal80*$^{ts}$ *x nrx-IV RNAI*$^{BL32424}$), n = 35 non-induced adult males and n = 55 induced adult males. *nrg* RNAi (*wor-GAL4, tub-Gal80*$^{ts}$ *x nrg RNAI*$^{BL37496}$), n = 35 non-induced adult males and n = 55 induced adult males. Data statistics: one-way ANOVA with a Kruskal–Wallis multiple comparison test. Results are presented as box and whisker plots.

Banach-Latapy et al, Supplemental Figure 1

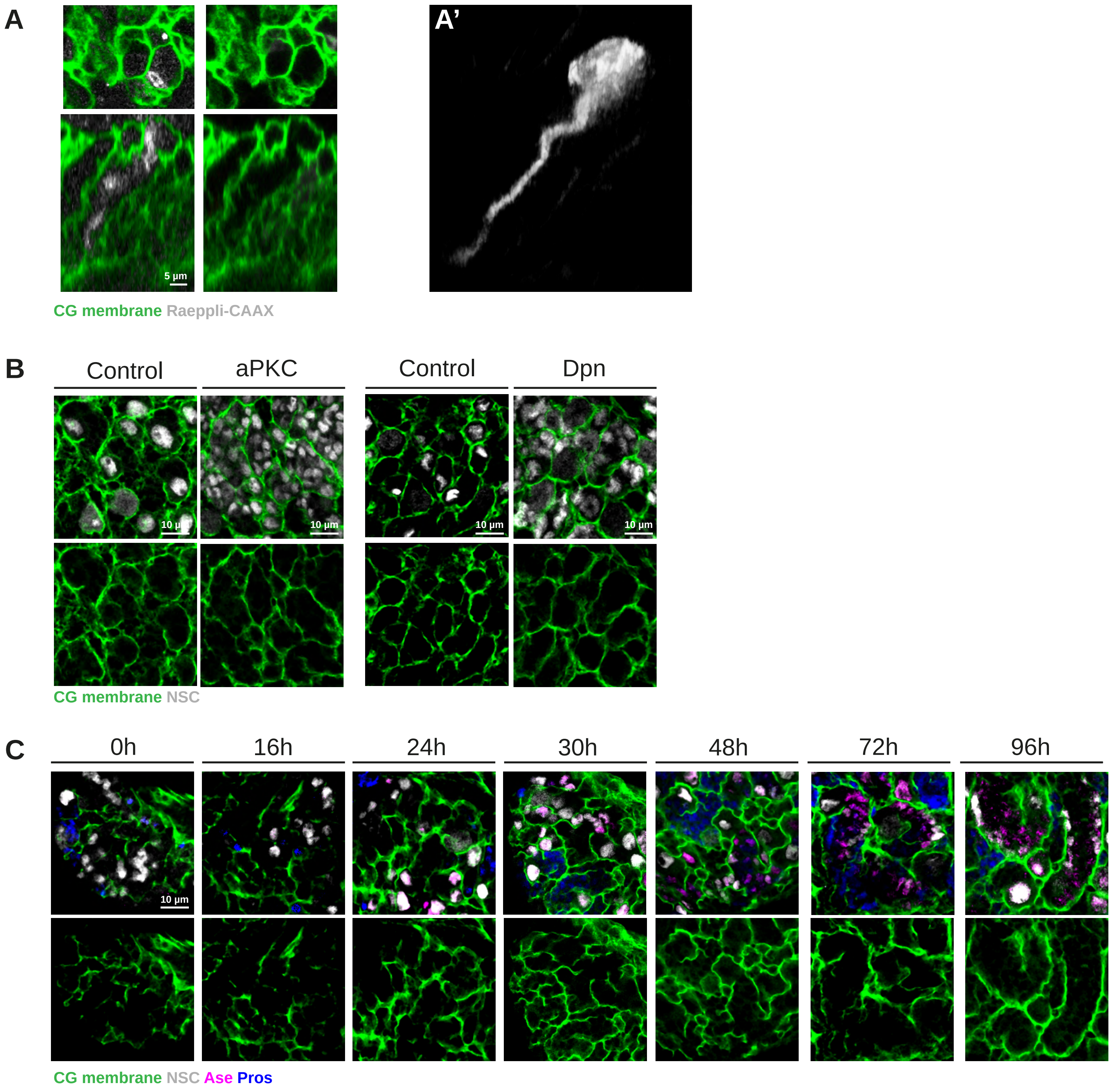

Banach-Latapy et al, Supplemental Figure 2

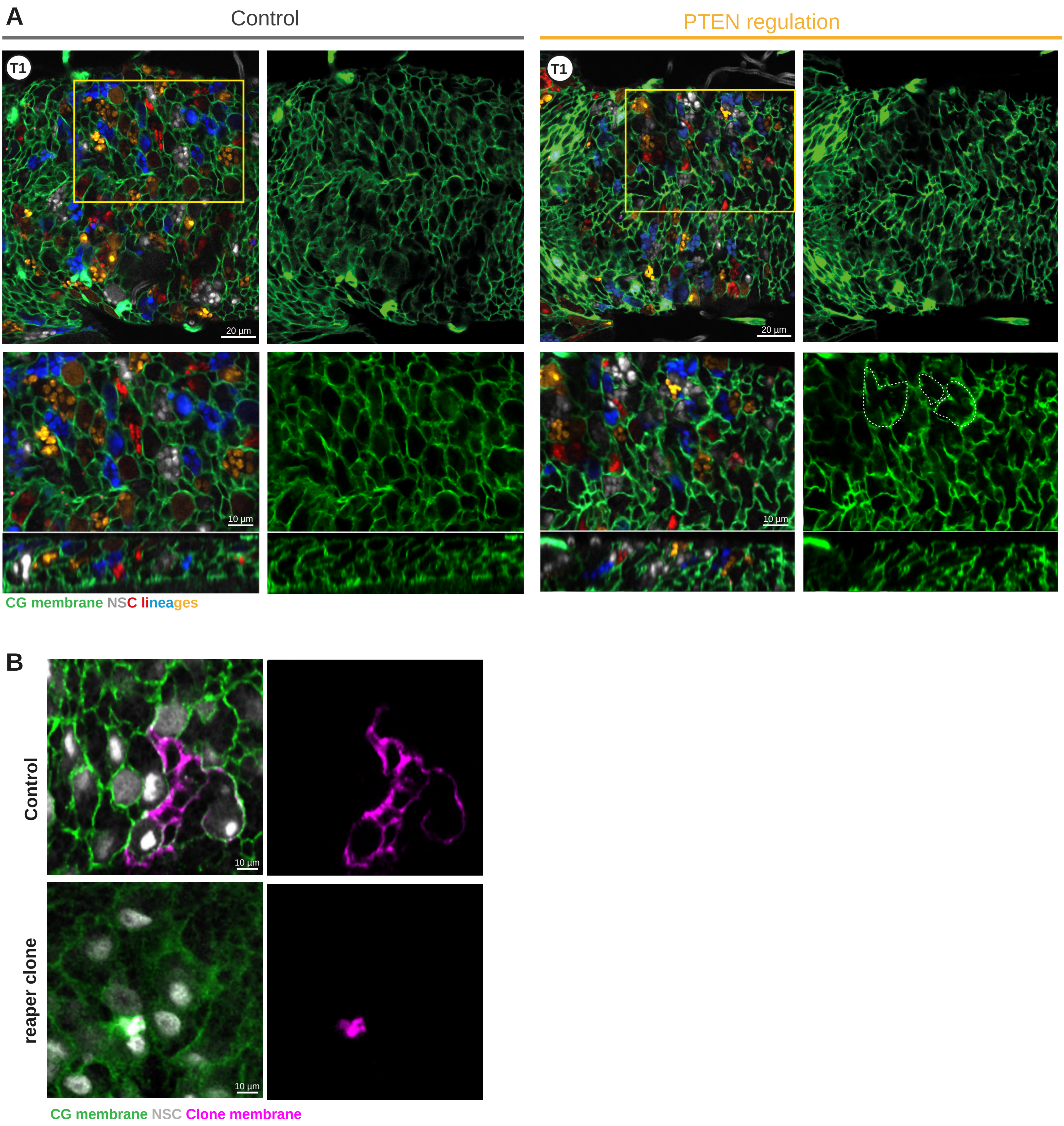

Banach-Latapy et al, Supplemental Figure 3

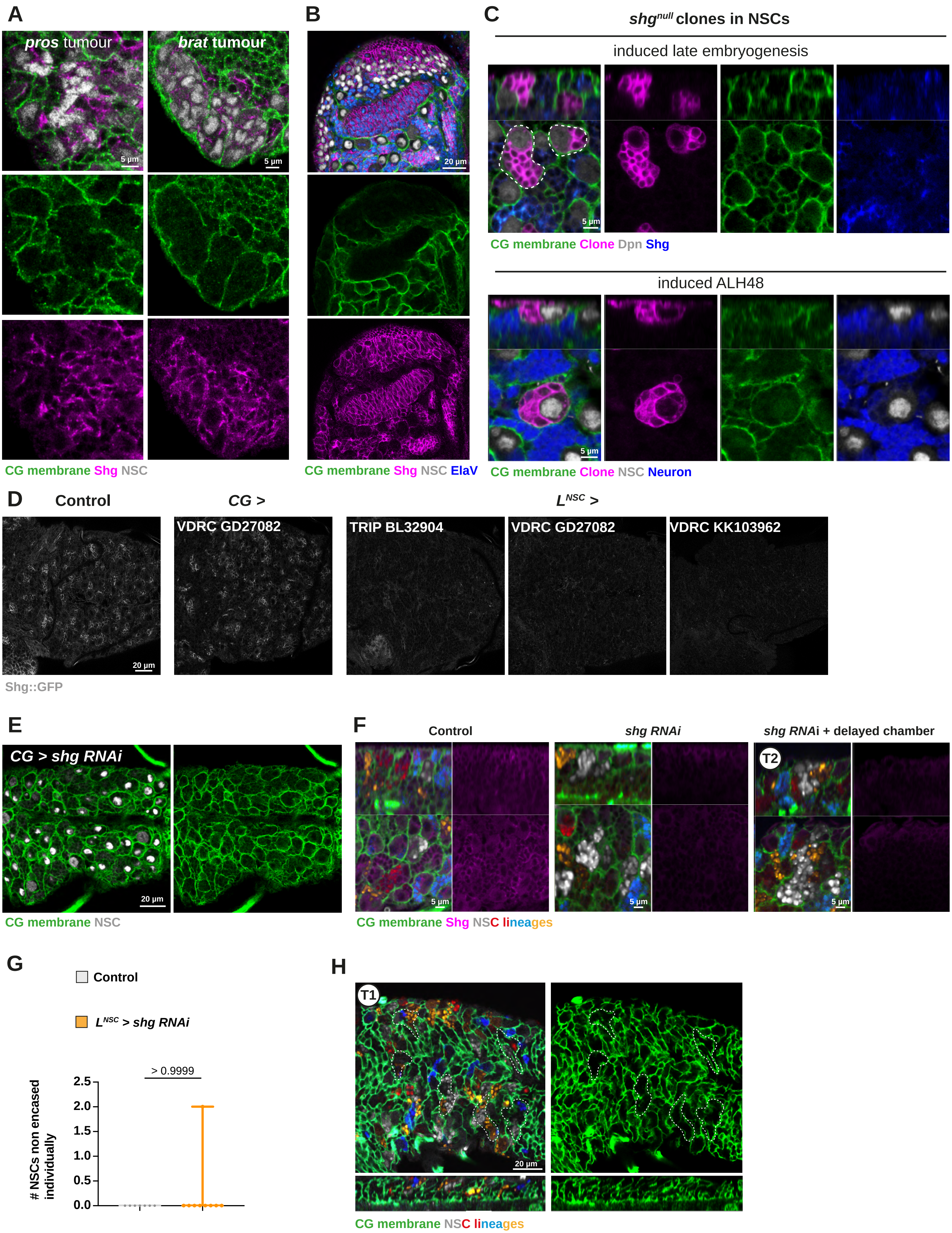

Banach-Latapy et al, Supplemental Figure 4

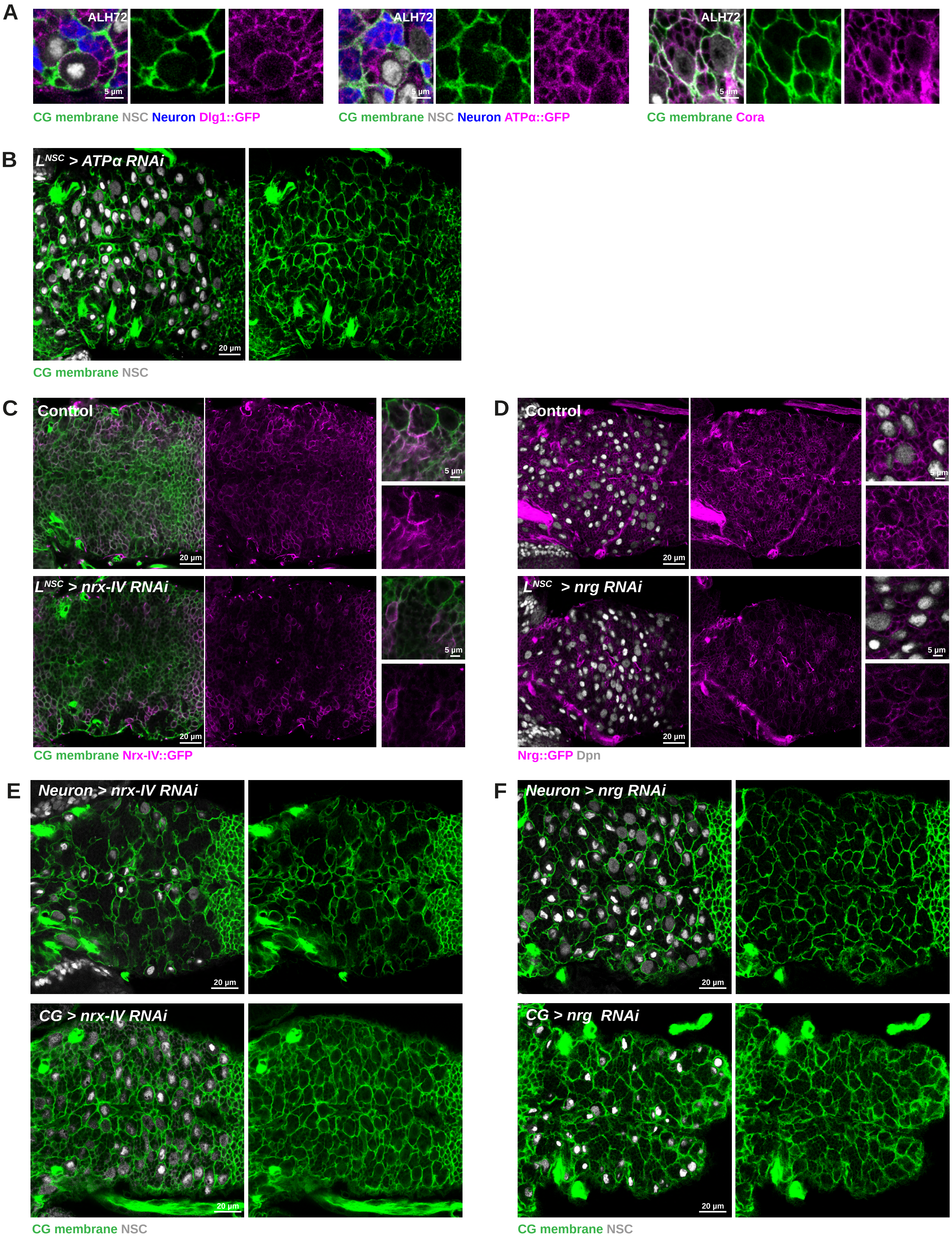

Banach-Latapy et al, Supplemental Figure 5

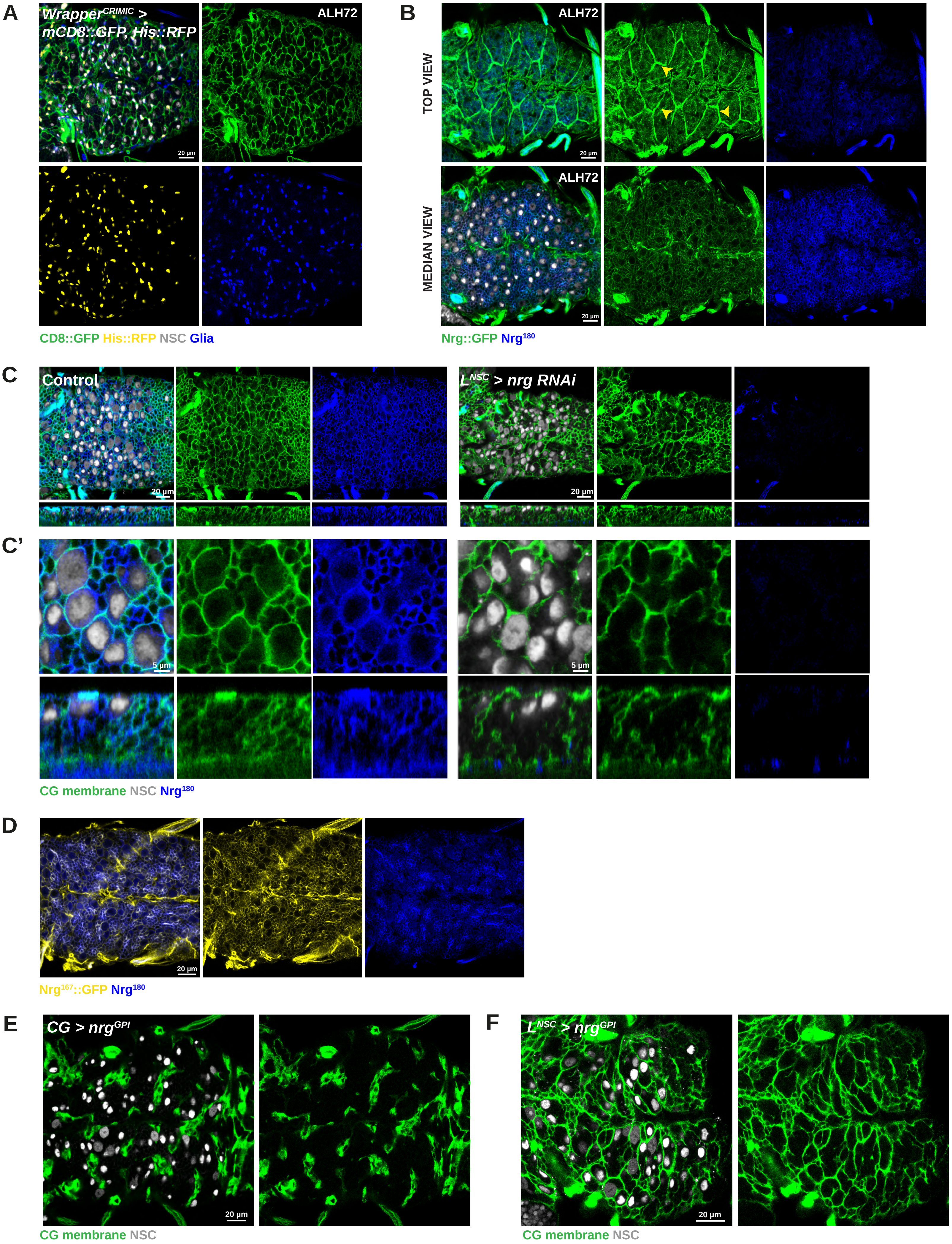

Banach-Latapy et al, Supplemental Figure 6

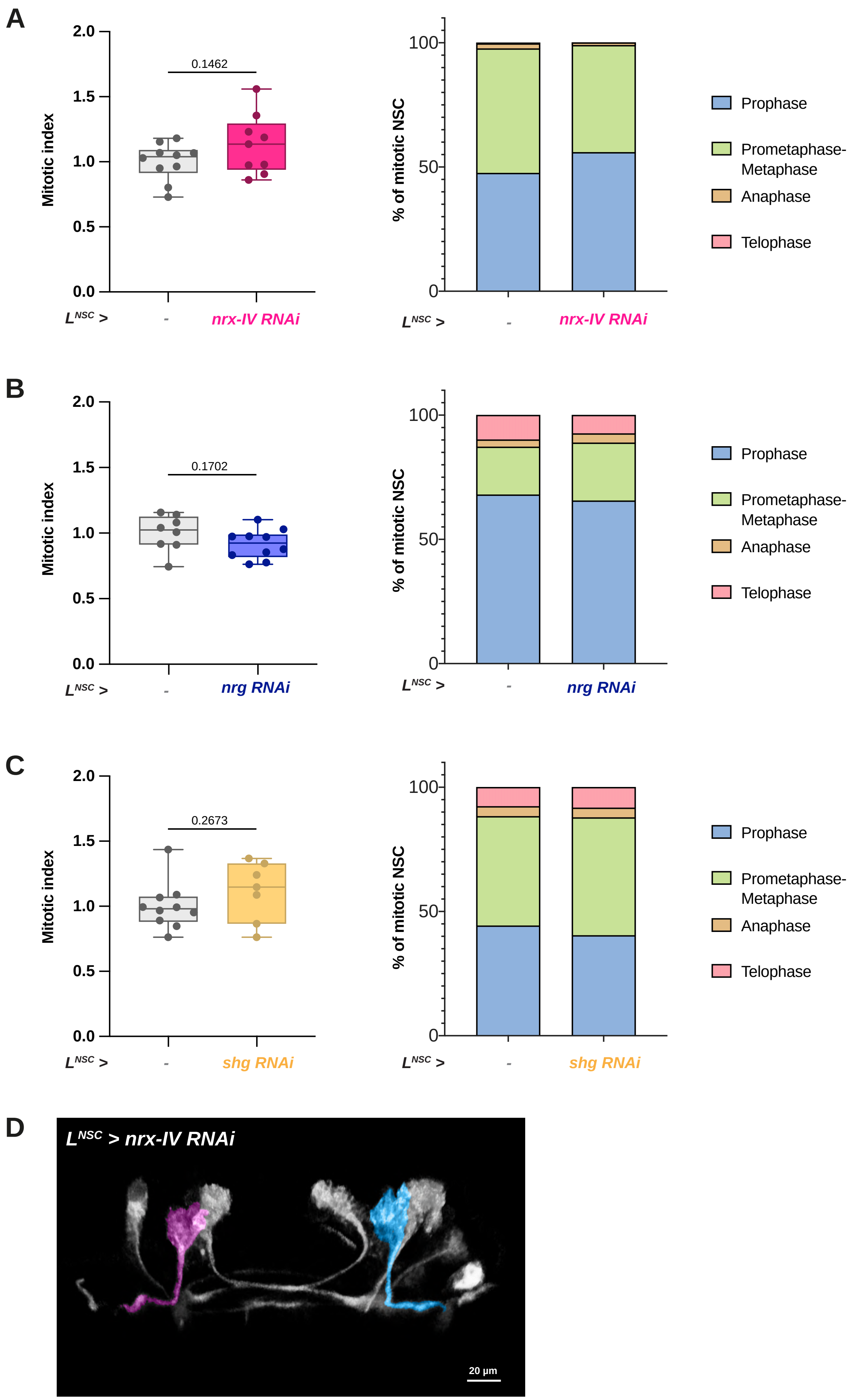

Banach-Latapy et al, Supplemental Figure 7

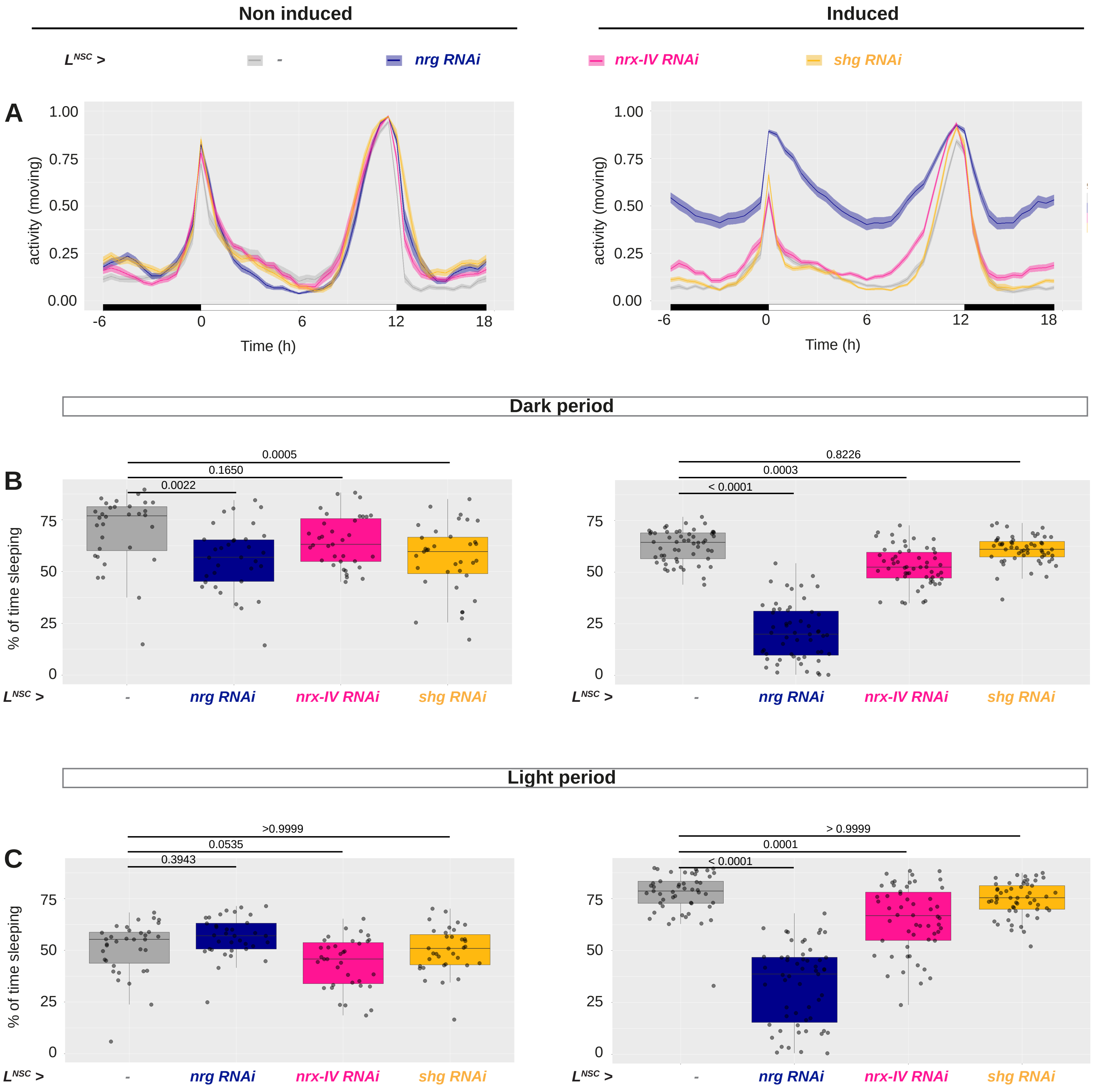
